# De novo design of antiviral monobody binders

**DOI:** 10.64898/2026.11.12.760042

**Authors:** Pedro Moreira, Shuhao Xiao, Carlota J.F. Conceição, Marta Alenquer, Diogo H. Silva, Mariana Parada, Madalena C. Marques, Joseph Schmidt, Carolina C. Buga, Christine Cruz-Oliveira, Jéssica Rodrigues, Marta Pires de Miranda, Beatriz Vieira-da-Silva, Ye Wei, Javier Collado, María Jesús Rodríguez-Espinosa, Rocío Arranz, Sandrine Georgeon, César Santiago, José María Valpuesta, Ana S. Veiga, Miguel A.R.B. Castanho, Isabel A. Abreu, Maria João Amorim, João B. Vicente, Bruno E. Correia, Cláudio M. Soares, Diana Lousa

## Abstract

Monobodies (Mobs) are a class of immunoglobulin (Ig)-like proteins that have emerged as promising therapeutic scaffolds for applications ranging from antiviral therapy to cancer treatment. Unlike conventional monoclonal antibodies (mAbs), Mobs can be readily expressed in *Escherichia coli* at high yields, offering important advantages in overall developability. However, discovering antigen-specific Mobs with high binding affinity traditionally requires screening large, randomized *in vitro* libraries. To optimize Mob discovery, we developed MobGen, a computational pipeline for *de novo* design of Mob binders against viral antigens. As a proof of concept, we generated Mobs that bind the SARS-CoV-2 Spike glycoprotein Receptor Binding Domain (RBD). Subsequent *in silico* optimization improved production yield, thermal and conformational stability, binding affinity, and antiviral activity, demonstrating a promising approach for rapidly developing antibody-like molecules that target viral entry. These findings highlight MobGen’s potential for *de novo* design of Mobs against emerging viruses with pandemic potential.

## INTRODUCTION

As a central component of host defense, the immune system deploys antibodies to defend against threats to its integrity. Refined over millions of years of host-pathogen co-evolution, antibody-mediated immunity can prevent pathogen entry and, when sterilizing immunity is not achieved, restrict infection and help reduce disease severity and onward transmission. These properties have made monoclonal antibodies (mAbs) an important class of biopharmaceuticals, offering high affinity and specificity toward defined molecular targets. However, their production generally relies on mammalian expression systems to support proper folding, disulfide-bond formation, assembly, and post-translational modifications (PTMs) (Rojekar et al., 2026), contributing to substantial manufacturing complexity and cost. To address these limitations, researchers have proposed several types of mAb-like molecules that may offer advantages over mAbs (Chaves et al., 2025a). Among these, monobodies (Mobs) have emerged as a promising alternative.

Mobs are synthetic monomeric proteins derived from the 10^th^ repeat of Human Fibronectin type III (FN3). They are usually 90 to 100 residues long, with a β-sandwich fold consisting of 7 β-strands, named A through G. This fold makes them similar to the variable regions of mAbs, with 3 of the Mob’s loops (BC, DE, and FG) acting as analogs of mAbs’ complementarity-determining regions (CDRs) that bind antigens (Koide et al., 1998). Compared to mAbs, Mobs are smaller proteins; they lack PTMs, and their stability does not depend on disulfide bonds. As a result, bacteria can produce Mobs at high yields, offering a notable economic advantage over mAb production. In addition, compared with nanobodies and minibinders generated by *de novo* design approaches, Mobs are less likely to be immunogenic because they are human-derived (Chandler and Buckle, 2020; Gebauer and Skerra, 2020; Koide and Koide, 2007).

Traditionally, Mobs are optimized in experimental settings by focusing on the CDR-equivalent BC/DE/FG loops to generate Mobs with mAb-like binding properties (Koide et al., 1998; Wojcik et al., 2010). However, it is also possible to mutate the CD loop (located on the C-terminal side) and the β-strands C and D to create a more structured interface (Koide et al., 2012). Researchers have engineered Mobs for oncologic diagnosis (Park et al., 2015; Ramakrishnan et al., 2019; Venu et al., 2024) and therapeutic applications (Grebien et al., 2011; Gupta et al., 2018; Spencer-Smith et al., 2016). Additionally, Mobs have been developed as inhibitors that block viral entry and neutralize viral infections by binding to relevant epitopes, predominantly viral host cell receptors. For example, in Kondo and colleagues’ work, phage display screened large in vitro libraries containing 3×10^13^ randomized sequences, enabling the identification of Mobs with picomolar binding affinity that target SARS-CoV-2 receptor-binding domains (RBD) and thus impair their ligation to the cellular receptor, angiotensin-converting enzyme 2 (ACE2) (Kondo et al., 2022, 2020). However, screening such large libraries to discover high-affinity binders is resource- and labor-intensive.

Recent advances in protein design enable the computational design of Mobs as protein binders (Eguchi et al., 2022; Perez et al., 2026). AF2-based (Evans et al., 2022; Jumper et al., 2021) design methods have shown significant promise across multiple protein design tasks, including miniprotein (Pacesa et al., 2025) and nanobody design (Mille-Fragoso et al., 2026; Swanson et al., 2025), achieving success rates unmatched by previous pipelines. Here, we propose MobGen, a Mob design pipeline that integrates AF backpropagation (Frank et al., 2024; Pacesa et al., 2025), the generative protein backbone method RFdiffusion (Watson et al., 2023), and ProteinMPNN (Dauparas et al., 2022) for *de novo* Mob binder design against a given input protein target structure.

As a proof of concept, we selected the D614G-containing SARS-CoV-2 spike protein, corresponding to Lineage B.1 (GISAID Clade G; Nextstrain Clade 20A), as our target. This variant provides a well-characterized model system with extensive structural and functional data. SARS-CoV-2 remains a relevant target because its continued evolution can produce antigenically distinct variants, sustaining the need for rapidly adaptable strategies to develop new binding molecules (Uzer et al., 2026). We generated over 10,000 sequences in total targeting the ACE2 binding motif of the SARS-CoV-2 RBD, from which ∼1,000 Mob sequences were selected for experimental validation. We used these sequences to build a tailored yeast display library (Chao et al., 2006; Uchański et al., 2019) and identified 7 SARS-CoV-2 binders, one of which showed viral-neutralizing activity with a micromolar IC_50_. We further optimized these binders *in silico* through sequence redesign with ProteinMPNN and scaffold engineering, yielding refined variants with improved stability and solubility, nanomolar binding affinities, and enhanced neutralization activity. Together, these results establish MobGen as an efficient computational pipeline for discovering and optimizing Mob binders.

## RESULTS

### MobGen: fold-conditioned design pipeline for Mob binder generation

Building on the recent success of the AF2-based “hallucination” approach for *de novo* binder design (Frank et al., 2024; Mille-Fragoso et al., 2026; Pacesa et al., 2025), we developed MobGen as a fold-conditioned design pipeline for Mob binders (**Figure 1A**). MobGen first leverages RFdiffusion to generate diverse starting poses for the Mobs with varied paratope length and conformation. AF2 subsequently co-folds the Mob and the target antigen, then designs the Mob to improve prediction confidence via backpropagation through the AF2 network. ProteinMPNN then redesigns the paratope. Finally, AF2 predicts the designs to select the highest-confidence candidates for experimental validation. A more detailed breakdown of the pipeline is provided in the **Methods** section.

**Figure 1:**
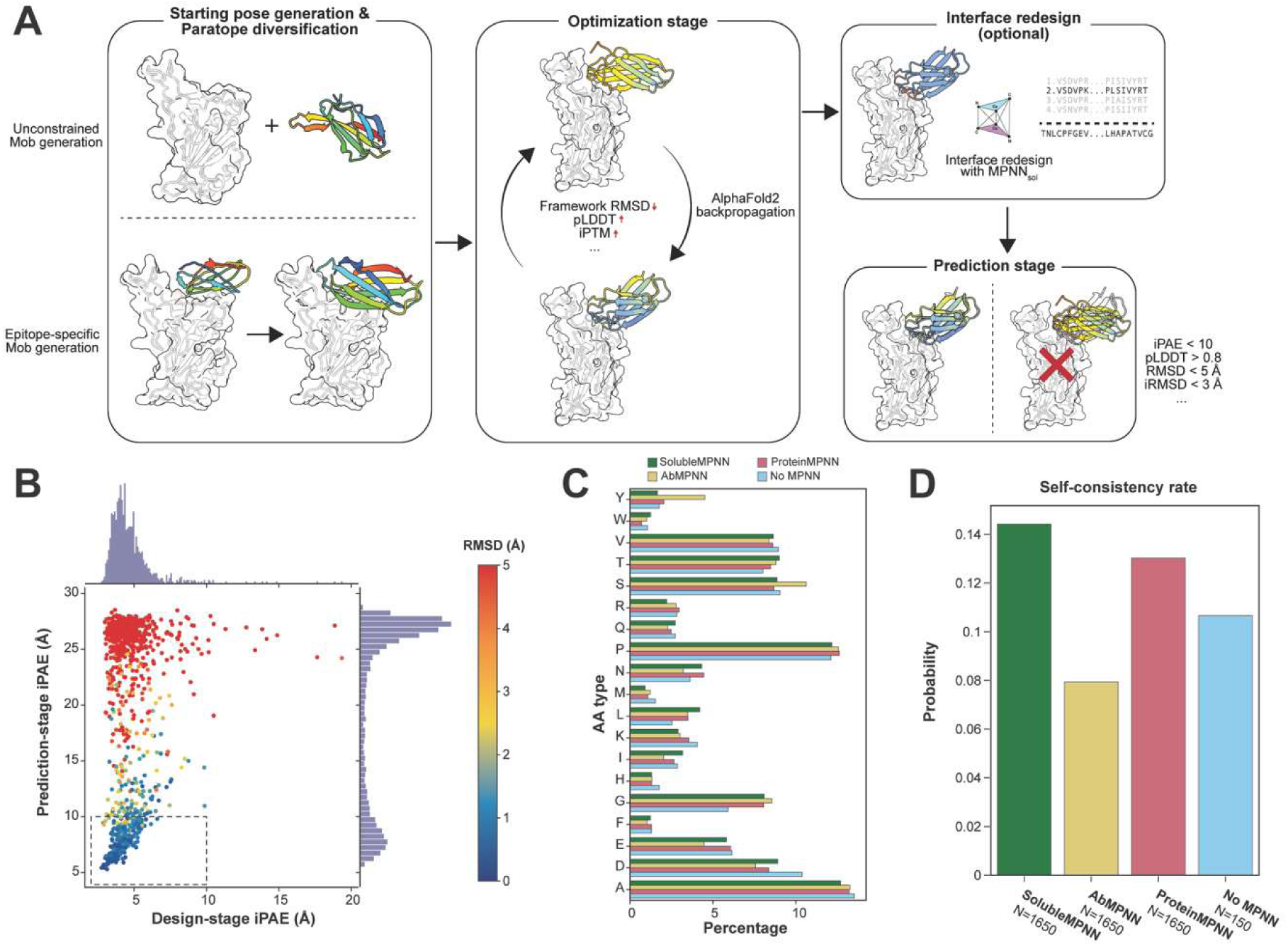
MobGen pipeline and evaluations of different ProteinMPNN models. (**A**) Main inference steps of the MobGen framework: RFdiffusion first generates starting Mob poses with diversified paratope lengths and conformations. Mobs are then designed via AF2 backpropagation, optimizing several confidence metrics; ProteinMPNN is then leveraged to further optimize the interface residues, particularly their solubility. Finally, designs are re-predicted with AF2 to check for self-consistency. (**B**) Relationship between the iPAE computed during the design stage and the prediction stage, with each design colored by the RMSD between the designed and predicted binder. Designs were generated with a SolubleMPNN loss. (**C**) Amino acid propensity in BC and FG loops when using different MPNN models for design, including designs generated without MPNN loss or redesign (No MPNN), for comparison. (**D**) Self-consistency rate (i.e., iPAE < 10 and RMSD < 2.5 Å) of the designs generated by different MPNN models.

Using MobGen to target the B.1 SARS-CoV-2 Spike RBD (PDB: 6M0J), we generated 11,450 sequences. We used interface PAE (iPAE) as the primary evaluation metric in both the design and prediction stages (**Figure 1B**) and accepted designs only if they demonstrated self-consistency (i.e., binder Root Mean Square Deviation (RMSD) between the design and prediction stages < 2.5 Å and pLDDT > 0.80). We first tested three ProteinMPNN model weights: the original ProteinMPNN (Dauparas et al., 2022), SolubleMPNN (Goverde et al., 2024), and AbMPNN (Dreyer et al., 2023). Although amino acid composition in Mob loops was broadly consistent across independently trained model weights, with AbMPNN showing a preference for tyrosine and serine (**Fig. 1C**), designs generated by different ProteinMPNN models varied in self-consistency (**Figure 1D**). SolubleMPNN achieved the highest *in silico* success rate and was therefore selected for interface refinement in the production campaigns.

In total, we selected 970 self-consistent designs to be screened using yeast surface display (YSD; **Figure 2A**). The initial screen showed a positive binding population, which was enriched after the second and third rounds of sorting (**Figure 2B** and **C**). After sorting, we identified 7 distinct sequences from the original *in silico* library’s binding population. Six of these proteins originated from a single binding pose (**Figure 2D** and **E**), while the remaining one came from a different backbone. During sequencing of the YSD results, we detected 9 additional binding sequences containing random mutations. As one of our aims is to discover proteins with potential antiviral activity, we decided to characterize these yeast-generated variants. But because these Mobs (Mobs 3560-3569) were not designed *in silico*, they should not be used in the validation of MobGen, and thus we did not include their data in the main body of the paper, only in the **Supplementary Data** section (**Tables S8** and **S9**, **Figures S19**, **S20** and **S21**).

**Figure 2:**
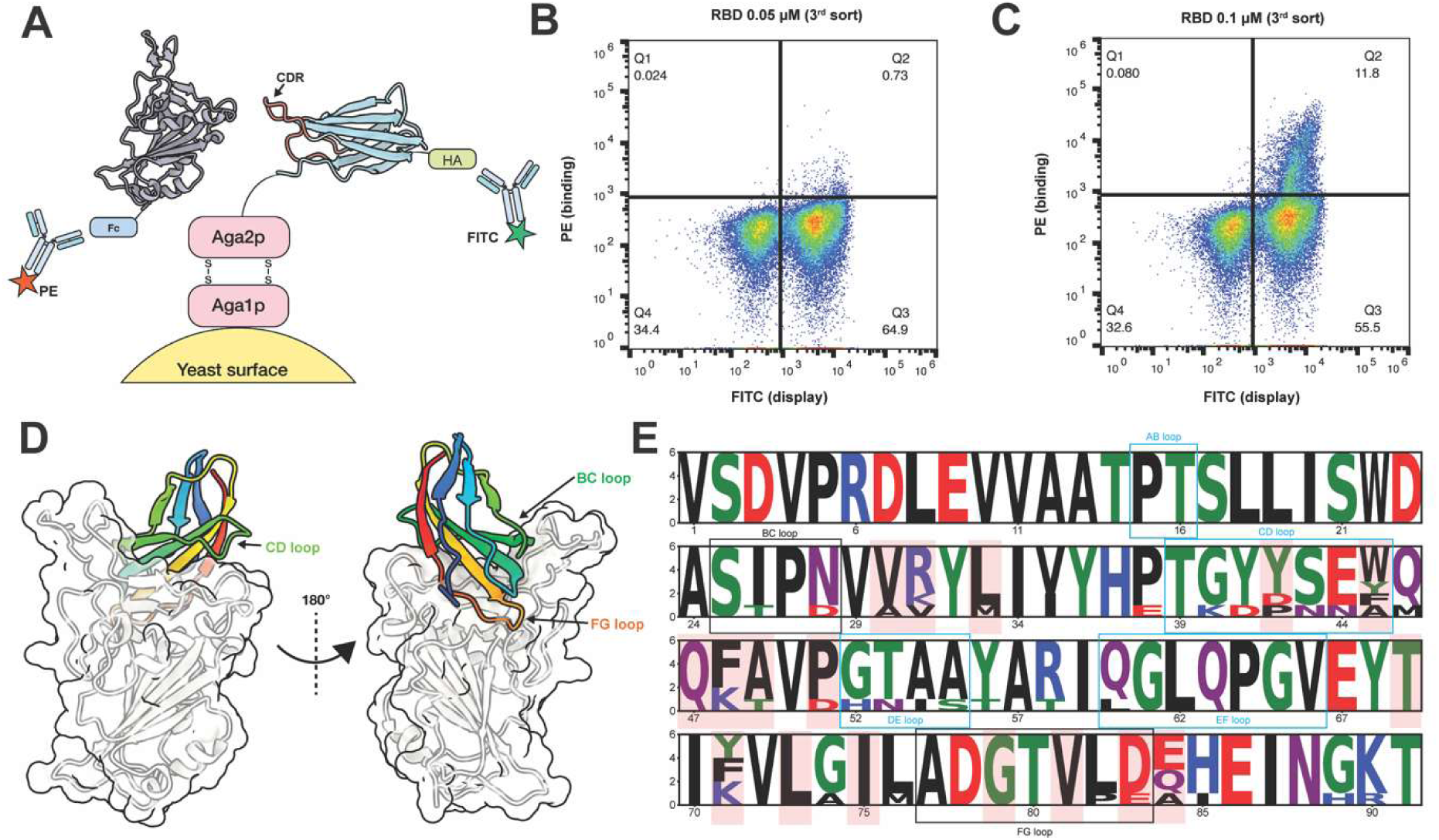
Yeast surface display results and predicted binding modes of the binders. (**A**) Schematic representation of the yeast display platform. (**B** and **C**) Representative flow plots of yeast display screening against the SARS-CoV-2 Spike RBD at different concentrations (50 and 100 nΜ). The x-axis shows Mob display (FITC signal), while the y-axis indicates RBD binding (PE signal). Double-positive populations for both Mob display and RBD binding appear in the upper-right quadrant. (**D**) Predicted structural representation of the binding pose shared by the 6 YSD-discovered binders (of the most common binding pose). (**E**) Sequence logo representation of the sequence diversity of the 6 YSD-discovered binders with the same binding mode.

When expressed and purified from bacteria, most of the YSD-screened binders could be produced only at modest yields (**Table S1**). They exhibited moderate thermal stability as analyzed by differential scanning fluorimetry (DSF), with melting temperatures (T_m_) in the 42.8-53.5 °C range, a single Mob having a higher T_m_ of 66.2 °C (**Table S1** and **Figure S1**). We also tested the neutralization activity of the library-identified Mobs against SARS-CoV-2 Spike WT (D614G)-pseudotyped lentiviruses. Among these candidates, Mob-2515 exhibited antiviral activity, with a half-maximal inhibitory concentration (IC₅₀) of 7.27 ± 0.54 μM (2 replicates; see **Figure S11** and **Table S5**).

These findings show that MobGen can generate promising Mob candidates that not only bind the selected target antigen identified through our initial Yeast Display screening but also inhibit viral entry. However, the suboptimal properties of the Mobs generated in this study underscore the need for further downstream optimizations to improve their therapeutic potential.

### *In silico* optimization improved thermal stability and binding affinity

First, we sought to improve the binders’ solubility, given their original low production yields and modest thermal stability (**Table S1** and **Figure S1**). Kondo and colleagues reported that adding a 7-residue C-terminal tail (amino acid sequence EIDKPSQ) increased the solubility of their candidate Mobs 3-fold (Kondo et al., 2022). We first performed Molecular Dynamics (MD) simulations of 2 strong-binding candidates from our initial library: Mob-2515 (with neutralizing capacity) and Mob-2550 (without detectable neutralizing capacity). We ran MD simulations of each with and without the C-terminal tail. We hypothesized that one key reason the tail can stabilize Mob could be hydrophobic interactions mediated by isoleucine at the second position of the tail sequence. Accordingly, we also performed MD simulations with only the first two tail residues (glutamic acid and isoleucine; referred to as “EI”).

We first predicted three versions of Mob-2515 and Mob-2550 using the AlphaFold3 (AF3) web server (Abramson et al., 2024): the original sequence without a tail (Mob-2515 and Mob-2550), with the first two residues of the tail (Mob-2515-EI and Mob-2550-EI), and with a full 7-residue tail (Mob-2515T and Mob-2550T). Importantly, these AF3 predictions showed consistent conformations across the three versions and matched the original MobGen designs. Based on these predictions, we ran 1 µs simulations with 3 replicates in GROMACS for each system (see **Methods**). Analyzing the temporal evolution of the entire Mob’s RMSD relative to the simulation’s initial position showed that, for Mob-2515, adding the tail did not clearly improve stability (**Figure S2**). However, the tail graft strongly affected Mob-2550, since the original protein had a significantly higher average RMSD (∼2.1 Å) than Mob-2550T and Mob-2550-EI (∼1.5 Å) (**Figure S2**). Visual inspection of the high-RMSD regions in Mob-2550 simulations shows a common feature: high fluctuation of the last β-strand (fragment G), which does not occur in Mob-2515 (**Figure 3A** and **B**). This pattern is also clearly seen in the Root Mean Square Fluctuation (RMSF) analysis (**Figure 3C**). In several frames, the last β-strand seems to detach from the rest of the Mob fold (**Figure 3D**). Adding the tail, even in its truncated form, greatly stabilized the β-strand in the simulations (**Figure 3E-F**), with the isoleucine in the tail likely contributing to the improved stability through hydrophobic contacts with up to two prolines in its spatial neighborhood (**Figure S3**).

**Figure 3:**
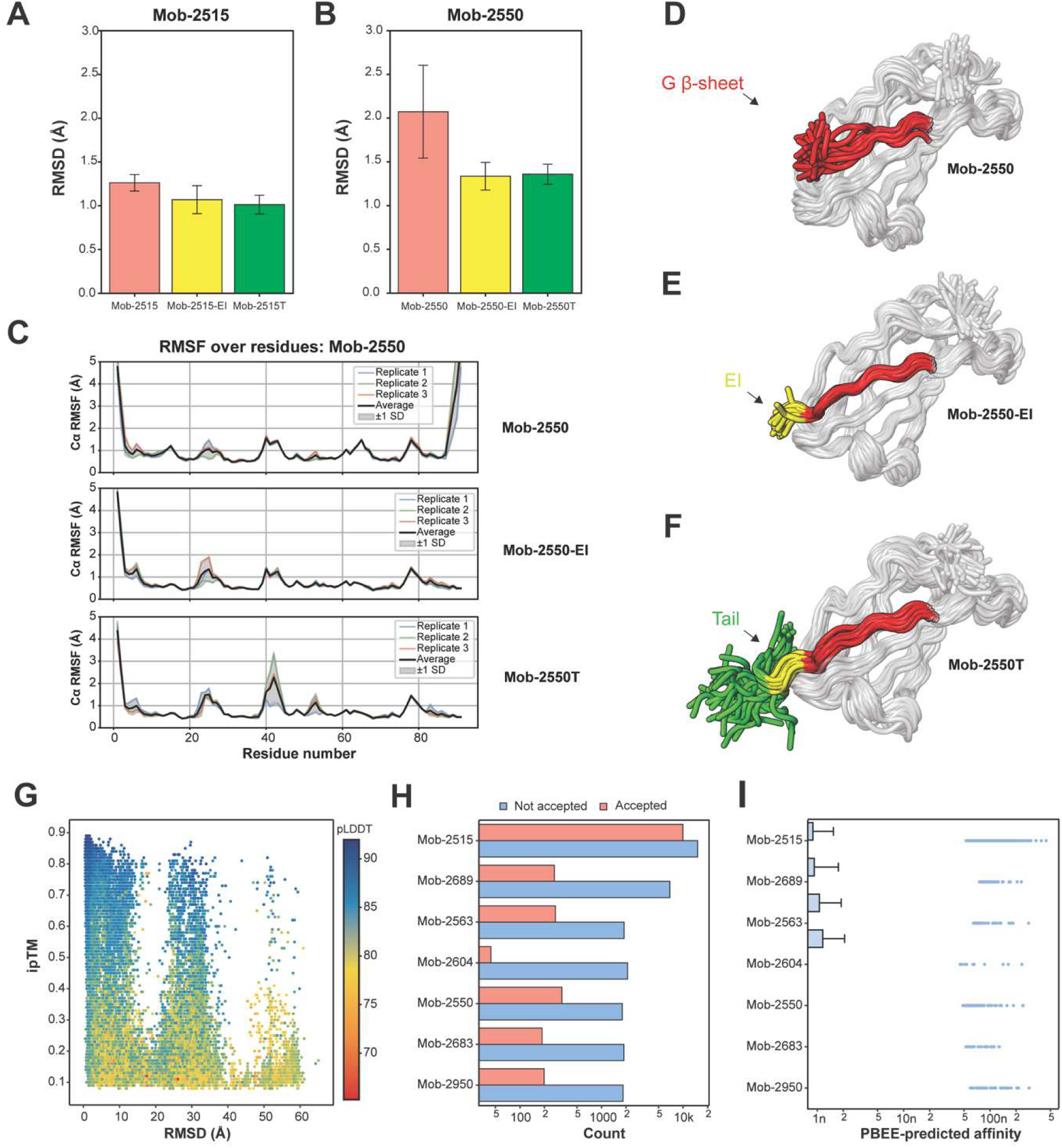
Effect of the C-terminal tail and sequence refinement with ProteinMPNN. (**A** and **B**) Bootstrap (95.45% confidence interval) of the RMSD of the G β-strand over the simulation time for the two Mobs, using 3 replicates for each simulation. (**C**) RMSF of the 1 μs simulation for Mob-2550, Mob-2550-EI, and Mob-2550T. To compare these Mobs of different sizes, we stopped the RMSF analysis at the last residue of the G β-strand. (**D-F**) Overlay of 50 frames from the MD simulation (each spaced 20ns apart) of a representative replica of Mob-2550. The red segment corresponds to the G β-strand (**D**); the yellow segment corresponds to the first two residues of that tail - EI (**E**); and the green segment corresponds to the rest of the tail (**F**). (**G**) AF3 metrics for the highest-confidence inference: RMSD between the design starting point and the mutant prediction versus ipTM, colored by complex pLDDT. (**H**) Count (on a log scale) of the accepted AF3 predictions, based on the design starting point. (**I**) Distribution of the PBEE-based affinity prediction of the AF3-accepted results.

Accordingly, we implemented this stabilization strategy by genetically fusing the 7-residue C-terminal tail to the other 5 YSD-discovered binders (Mob IDs 2563, 2604, 2683, 2689, and 2950) and the two simulated Mobs (2515 and 2550). As intended, these engineered YSD-discovered binders showed a clear improvement in expression yield, compared with the original Mobs, under the same expression conditions, with an average 7-fold increase in production, ranging from 2.6-fold for Mob-2515T to 11.4-fold for Mob-2950T (**Figure 4B**; **Table S1**). The production yield may corroborate the MD simulation prediction, where G β-strand stabilization is more pronounced in Mob-2550T (which showed a 10.8-fold yield improvement) than in Mob-2515T. We also observed that introducing the tail improved thermal resistance for several Mobs (**Figures 4C**, **S1** and **Table S1**). Further, dynamic light scattering (DLS) characterization showed that the designed Mobs had the expected hydrodynamic radii of monomeric Mobs (**Figure S6**) and were overall monodisperse. Far-UV circular dichroism (CD) spectra (**Figure S8**) exhibited the predicted predominantly β-sheet secondary structure. Some of the spectra displayed a positive band centered at 225-230 nm which, in the absence of any possible disulfide bridge, has been attributed to the clustering, relative positions and solvent exposure of tryptophan, tyrosine and phenylalanine residues in various proteins (Alston et al., 2004; Grishina and Woody, 1994; Woody, 1994).

**Figure 4:**
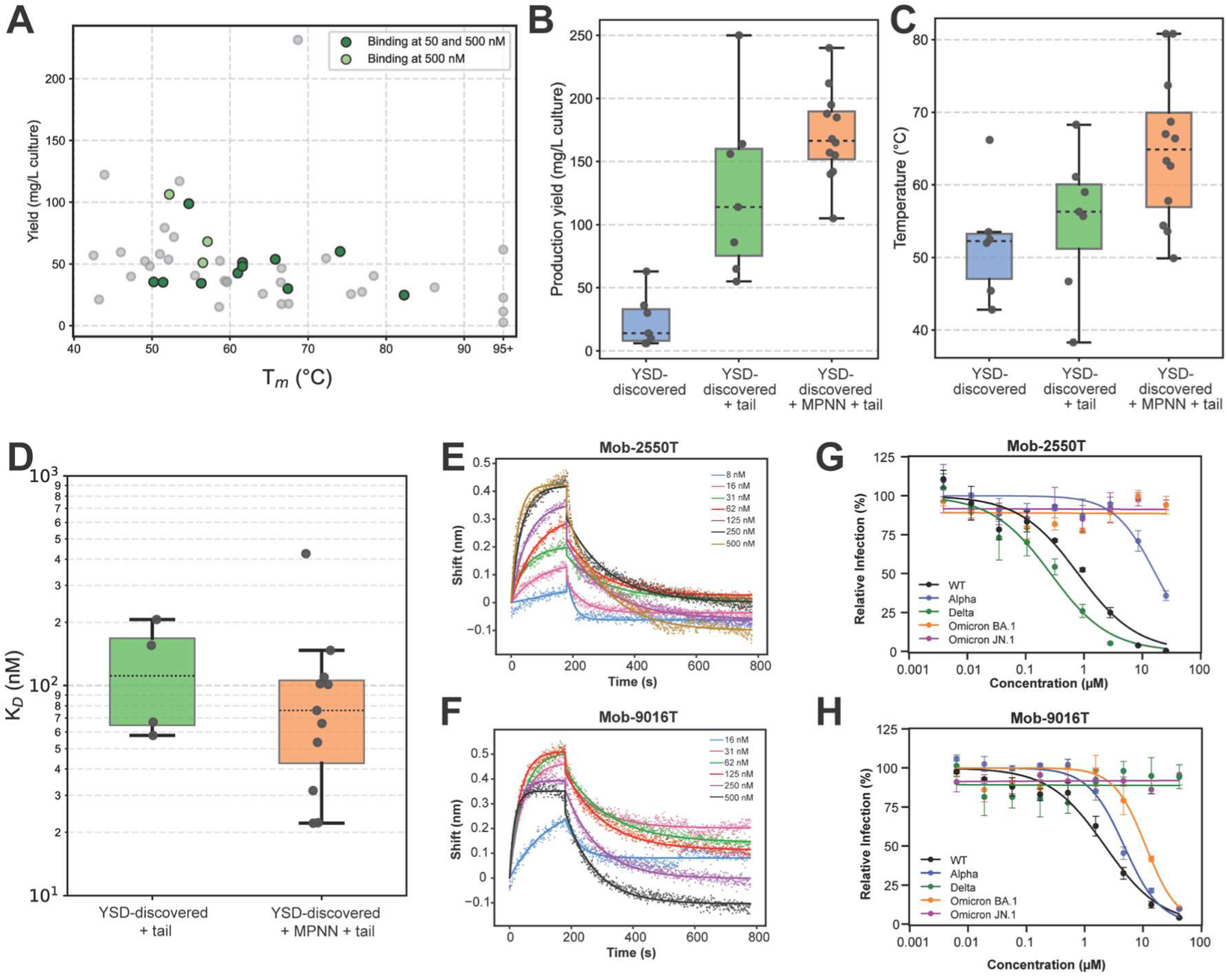
Experimental validation of YSD-discovered and *in silico* optimized Mobs. (**A**) Production yield and melting temperature measured by the mid-throughput platform for the ProteinMPNN-refined Mobs, each design colored by the binding signals detected by SPR at two RBD concentrations (50 and 500 nM) (**Figure S4**). (**B**) Comparison of the production yields in *E. coli* and (**C**) the melting temperature (T_m_) of the Mobs identified in the original YDS (blue), the same Mobs with the C-terminal tail (green), and their ProteinMPNN-refined designs with the C-terminal tail (orange). (**D**) Comparison of binding affinities (KD; log scale) between the Mobs identified in the original library after adding the C-terminal tail (green) and the ProteinMPNN-refined designs (orange). (**E** and **F**) Binding curves of Mob-2550T (E) and Mob-9016T (F) measured by SPR. (**G** and **H**) Neutralization activity of Mob-2550T (G) and Mob-9016T (H) against pseudoviruses across multiple SARS-CoV-2 strains.

We then used ProteinMPNN to redesign the initial 7 binders, using their predicted AF2 complexes from MobGen as starting points. In total, 42,697 non-redundant sequences were generated from these 7 binders. These sequences were subjected to AF3 predictions (Abramson et al., 2024) to test for self-consistency (**Figure 3G** and **H**) and PBEE (Chaves et al., 2025b), an ensemble model that leverages Rosetta-based biophysical metrics (Leaver-Fay et al., 2011) to predict binding affinity (**Figure 3I**; see **Methods**). We selected the top 50 refined Mobs, all bearing the aforementioned C-terminal tail, for screening on a custom-designed mid-throughput platform that enables screening up to 96 proteins per round, including bacterial expression, cell lysis, affinity chromatography purification, thermal stability, and target-binding assessment (**Figure 4A**; see **Methods**). This platform enabled us to test the production yield, melting temperature, and binding of 50 Mobs simultaneously in under 2 weeks (**Table S2** and **Figure S4**). Based on experimental filtering focused on apparent binding potency, we identified 12 Mobs with improved properties across assays.

After scaling up production and purification, the refined Mobs produced yields comparable to the YSD-discovered Mobs after adding the C-terminal tail (171 ± 36 and 127 ± 69 mg/L of culture; **Figure 4B; Tables S1** and **S3)**. Sequence redesign improved thermal stability in several Mobs; three of these (Mob-9007T, 9030T, and 9044T) displayed Tm values above 70 °C (**Table S3**), higher than those identified in the initial library screening (**Figures 4C**, **S1**, **S5** and **Tables S1 & S5**). In these Mobs, we also observe similar patterns in the DLS data (expected hydrodynamic radii and essentially monodisperse preparations; **Figure S7**) and CD analyses (β-sheet secondary structure with a positive band at ∼225-230 nm; **Figure S9**) compared with the YSD-discovered Mobs with the added tail. By fluidics-free surface plasmon resonance (SPR) assays, we analyzed the binding kinetics for 15 Mob binders, all bearing the C-terminal tail (2515T, 2550T, 2563T and 2683T from the initial library and 11 from the ProteinMPNN-refined Mobs), determining average K_D_ values ranging from 22.1 to 424.8 nM (**Figure 4D** and **S10**; **Table S4**). ProteinMPNN refinement had little impact on binding affinity but affected thermal and conformational stability (**Figure 4B, C**, and **D**).

Thus, this two-step refinement (adding the C-terminal tail and ProteinMPNN refinement) improved the thermal stability of the Mob binders, increasing their value as potential biopharmaceuticals.

### *De novo-*designed Mobs neutralize SARS-CoV-2

We then tested the neutralization activity of these *in silico*-refined Mobs against SARS-CoV-2. For designs showing RBD binding by SPR, we evaluated antiviral activity against both spike B.1-pseudotyped lentiviral particles and SARS-CoV-2 B.1 authentic virus. Two Mobs from the original YSD screening, Mob-2550T and Mob-2563T, engineered to bear the C-terminal tail, showed neutralization activity in both assays, with averaged IC_50_ values below 10 μM (**Figures S12 & S13, Table S6**). Both neutralization assays identified Mob-2550T as the Mob with the lowest IC_50_ values: 1.49 ± 0.15 μM against pseudotyped viruses and 3.2 ± 0.4 μM against SARS-CoV-2 B.1 (**Figures 4E, S12 & S13, Table S6**). Not every effort to improve properties increased neutralizing activity. In fact, while Mob-2515 showed the highest neutralizing activity against the pseudotyped viruses (7.27 ± 0.54 μM) among the pre-refinement Mobs, Mob-2515T showed an increased IC_50_ value of 24.42 ± 3.73 μM, indicating that the addition of the C-terminal tail is likely detrimental to its antiviral activity (**Figures S12 & S13, Table S6**).

Among ProteinMPNN-refined Mobs, all platform-selected binders showed neutralizing activity (in the pseudovirus assays), with IC_50_ values ranging from ∼2 to ∼25 µM (**Figures S12 & S13, Table S6**), indicating little to no improvement in neutralization capacity despite the increased hit rate. Mob-9011T and 9016T showed IC_50_ values of 2.08 ± 0.28 and 2.40 ± 0.48 μM against pseudovirus and 8.8 ± 1.0 and 8.5 ± 1.4 μM against SARS-CoV-2 B.1, respectively (**Figures 4F, S12 & S13, Table S6**).

To assess the breadth of the most potent Mobs described above (Mobs 2550T, 2563T, 9011T, and 9016T) against different SARS-CoV-2 variants (**Figures 4 G** & **H** and **S14**, **Table S7**), we performed neutralization assays with pseudotyped viruses carrying the Spike protein sequences corresponding to the Alpha, Delta, Omicron BA.1, and Omicron JN.1 variants. We observed that Mob-2550T displays antiviral activity against Alpha (IC_50_ = 16.7 ± 3.0 μM) and Delta (IC_50_ = 0.267 ± 0.051 μM), but does not neutralize the two tested Omicron variants (BA.1 and JN.1) (**Figure 4G**). Meanwhile, Mob-9016T also neutralizes Alpha (IC_50_ = 4.75 ± 0.38 μM) and Omicron BA.1 (IC_50_ = 11.16 ± 1.54 μM), but shows no activity against Delta or Omicron JN.1 (**Figure 4H**). Among the other tested Mobs, Mob-2563T also neutralizes the Delta variant but is inactive against the other tested variants, while Mob-9011T neutralizes only the Alpha variant among the variants analyzed (**Figure S14, Table S7**).

Overall, these results show that individual Mobs retain measurable antiviral activity against different SARS-CoV-2 variants, but neutralization breadth differs between Mobs, with each displaying a distinct pattern of variant susceptibility and escape.

### Molecular basis of the neutralization activity

To further investigate the binding interfaces of the most potent Mobs using MD simulations and MMPBSA calculations (**Figure 5A** and **B**), we simulated the Mob with the lowest averaged IC_50_ (Mob-2550T) and the refined Mob combining the lowest IC_50_ and highest T_m_ (Mob-9016T). Both Mobs are predicted to compete with the ACE2-binding domain (PDB: 6M0J; **Figure 5C**), with their binding modes favoring hydrophobic interactions. This is not surprising given the chemical characteristics of the RBD-ACE2 interface. However, several other interactions also occur, as indicated by the MMPBSA contributions from MD simulations of both complexes (3 replicates for 1 μs; **Figure 5A** and **B**).

**Figure 5.**
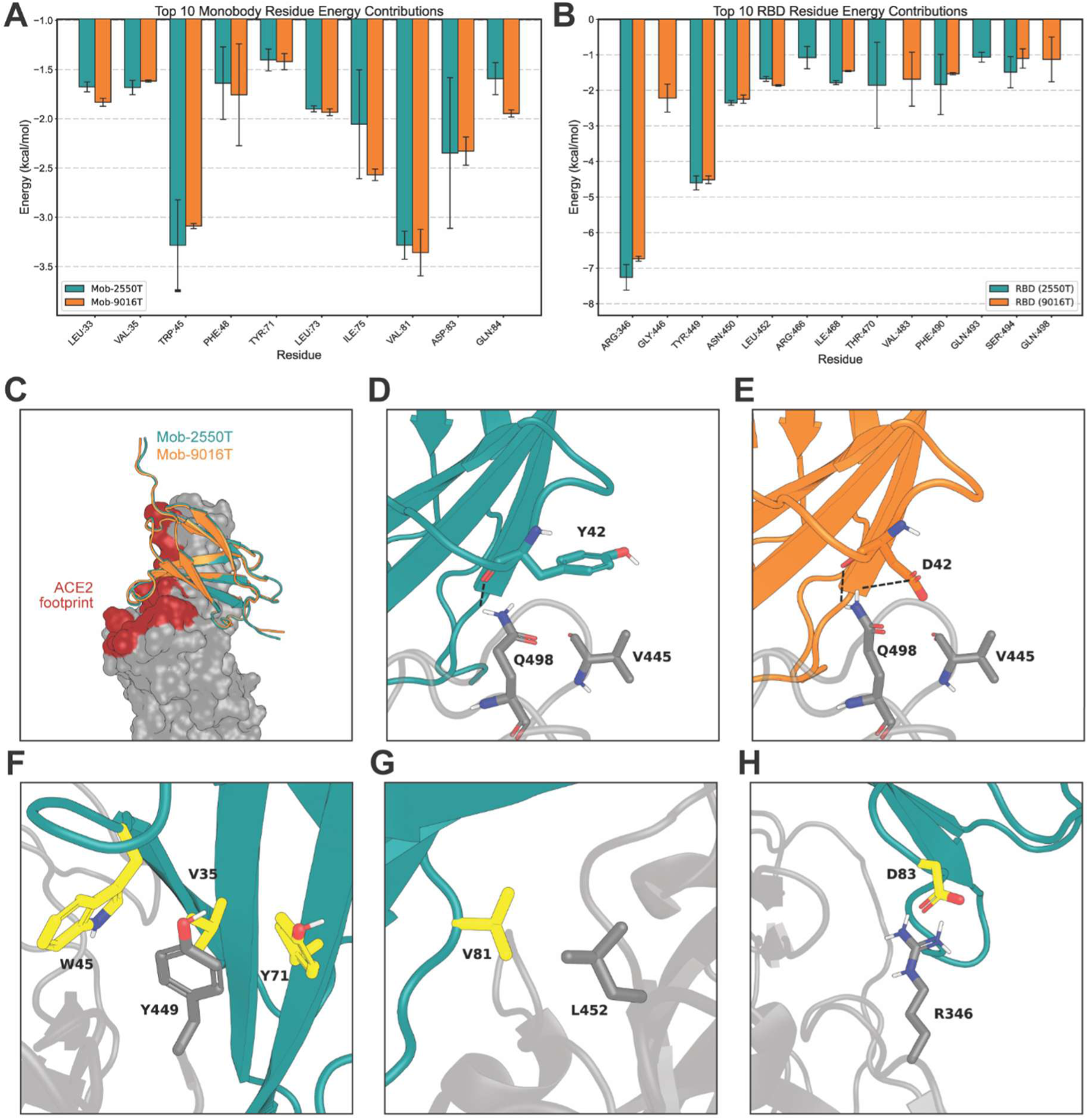
MMPBSA decomposition of the top 10 residue contributors to the interface energy of the selected Mobs. (**A** and **B**) MMPBSA binding energy decomposition of the last 500 ns of each of the three replicates of the MD simulations for Mob-2550T and Mob-9016T, in complex with the RBD. The top 10 residues by energy contribution per protein (A). The top 10 contributing RBD residues from the same analysis for the simulations (B). (**C**) Overlay of the ACE2 footprint (red) and the binding mode of Mob-2550T (teal) and Mob-9016T (orange) on the RBD. (**D** and **E**) Different interface residues between Mob-2550T (teal) and Mob-9016T (orange), with the promoted hydrogen bonds marked as dashed lines. (**F-H**) Shared interacting residues (yellow) of both Mobs that are predicted to be the highest contributors in terms of binding energy by the MMPBSA analysis.

These two Mobs differ by three residues (positions 31, 42, and 76), and only one mutation significantly increases interaction contributions at the binder-target interface in the MMPBSA analysis. In Mob-2550T, Y42 promotes hydrophobic interactions with RBD V445 and forms a hydrogen bond with the Q498 backbone (**Figure 5D**). In contrast, Mob-9016T loses this hydrophobic interaction because it has an aspartic acid instead of a tyrosine at position 42, but gains an additional hydrogen bond with Q498’s side chain in the RBD (**Figure 5E**). This additional interaction is also reflected in the minor contribution of RBD Q498 in the MMPBSA analysis (**Figure 5B**). According to this analysis, some of the most critical interactions on both Mobs include π-stacking interactions promoted by W45 and Y71 of both Mobs with Y449 of the RBD, coupled by additional nonpolar contacts to V35 of both Mobs (**Figure 5F**); hydrophobic contacts between V81 of both Mobs and L452 of the RBD (**Figure 5G**); and a salt bridge promoted by D83 of both Mobs to R346 of the RBD (**Figure 5H**).

The simulations also provided plausible explanations for the significant differences in K_D_ and IC_50_ values. Throughout the simulation, both Mobs most often interacted with five or fewer RBD residues (**Figure S15**), suggesting relatively modest overlap in their binding interfaces. Nevertheless, the Mob-2550T:RBD and Mob-9016T:RBD complexes remained stable over the simulation, as indicated by Mob RMSD values below 2 Å relative to their starting position (**Figure S16**) and low Mob fluctuation (RMSF < 1.5 Å, excluding the termini) (**Figure S17**). These observations suggest that the Mobs can establish stable binding interactions with the RBD. Moreover, both Mobs competed with ACE2, as verified by competition assays (**Figure S18**). Therefore, despite a predicted stable RBD-binding interface and measurable ACE2 competition, the small interface overlap likely limits their effectiveness as antiviral proteins.

## DISCUSSION

In this work, we developed MobGen, a *de novo* design framework for generating Mob binders. MobGen combines RFdiffusion, which generates starting points with a Mob-like fold and diversified loop lengths and conformations, and AF2, which designs Mob sequences via neural-network backpropagation. Additionally, ProteinMPNN is leveraged to refine the Mob paratope’s affinity and solubility properties. As a proof of concept, we targeted the SARS-CoV-2 spike protein from the lineage B.1 at the RBD site and developed neutralizing Mobs against this virus. Our computational pipeline generated promising candidates against the viral fusion protein target, requiring a library with up to 10 orders of magnitude fewer sequences than traditional experimental screening-based approaches (**Figure 2B**). From this initial library, we identified one generated Mob that displayed antiviral activity, with an IC_50_ in the single-digit micromolar range against the B.1 variant (**Figure S11** and **Table S5**).

We then applied two optimization strategies: adding a C-terminal tail and redesigning the sequence using ProteinMPNN. We combined AF3 confidence metrics with a biophysics-based affinity predictor to select candidates, prioritizing sequences supported by these complementary approaches. These interventions improved expression yield and thermal stability for several Mobs (**Figures 4B** and **C**), while selected variants displayed RBD-binding affinities in the low-nanomolar to submicromolar range (**Figure 4D**). The improvements were nevertheless design-dependent: the C-terminal tail did not improve every property of every Mob, and ProteinMPNN refinement did not consistently improve binding affinity relative to the tail-containing MobGen designs.

The limited effect of ProteinMPNN refinement on affinity may reflect the relatively small number of mutations introduced at the binding interface. Alternatively, MobGen-generated interfaces may already have been close to a local optimum within the sampled sequence space, despite less favorable developability properties. Distinguishing between these possibilities will require a more systematic comparison of alternative redesign strategies and direct affinity measurements for matched variants before and after refinement.

SPR-measured binding affinity did not show an obvious relationship with neutralization potency. This discrepancy may arise partly from the relatively rapid dissociation rates of the Mobs, which could limit target occupancy under the conditions used in the neutralization assays. In addition, the predicted Mob epitopes overlap only partially with the ACE2-binding footprint, potentially limiting steric inhibition of receptor engagement. Consistent with this interpretation, the Mobs competed with ACE2 (**Figure S18**) but showed only micromolar neutralization potency (**Figures S12 & S13, Table S6**). Some designs nevertheless retained activity against more than one SARS-CoV-2 variant, although none neutralized all variants tested (**Figures 4G, 4H** and **S14**; **Table S9**). These results indicate cross-variant activity, highlighting the potential therapeutic value of these *de novo* binders.

In the present implementation of MobGen, no explicit objective was used to discourage interactions mediated by the FN3 framework. Consequently, most designs were predicted to bind through the β-sheet surface rather than primarily through the BC, DE, and FG loops. Such structured interfaces may be easier for AlphaFold-based methods to model accurately. Framework-mediated binding has previously been demonstrated for Mobs (Koide et al., 2012), but it generally requires extensive mutation of the FN3 scaffold. Accordingly, the final neutralizing Mobs retained only approximately 26% sequence identity to the parental FN3 domain. This extensive sequence divergence weakens the assumption that these proteins will retain the low immunogenicity commonly associated with human-derived Mob scaffolds.

To address this limitation, we added objectives that preserve sequence identity to a selected FN3 scaffold and favor target contacts through the paratope loops. However, the lower accuracy of AlphaFold-based methods for loop-mediated interfaces may limit both the generation and reliable computational selection of such designs (Hitawala and Gray, 2025; Perez et al., 2026). Experimental evaluation of these alternative design modes is needed to determine whether they improve scaffold conservation without compromising binding success.

Further development of MobGen should therefore focus on several priorities. Because these designs diverge extensively from the parental human FN3 sequence, immunogenicity risk should be evaluated directly, while cytotoxicity and other safety properties should be assessed independently. Broader specificity and off-target binding studies will also be required before the designs can be considered therapeutic candidates. Additional opportunities include redesigning the Mobs against antigenically divergent viral variants and combining binders that recognize non-overlapping RBD epitopes. Multivalent or multispecific formats incorporating a Mob with greater overlap with the ACE2 footprint may improve neutralization potency and reduce susceptibility to viral escape.

The design framework could also benefit from newer complex-prediction and generative architectures. AF3-related approaches, Chai-1 (Chai Discovery et al., 2024), and Boltz-2 (Passaro et al., 2025) may provide improved models for structure-based sequence optimization and candidate filtering. Generative frameworks such as BoltzGen (Stark et al., 2026) may offer an alternative route to designing binder backbones and interfaces. However, their value for Mob design will need to be established through prospective experimental benchmarking rather than confidence metrics alone.

Overall, this study provides proof of concept that computational design can generate monobody proteins that bind a specified viral target and display measurable antiviral activity. The results also identify key limitations—including modest neutralization potency, incomplete variant coverage, extensive scaffold divergence, and inconsistent gains from sequence refinement—that define priorities for further MobGen development.

## METHODS

### MobGen Framework

For Mob generation, we used an FN3 structure (PDB: 1TTG) (Leahy et al., 1992) as the starting-structure template. The CDR-equivalent loop conformations and the template Mob length can be randomized using RFdiffusion. Mobs in the literature (Koide et al., 2012; Kondo et al., 2020) inspired the number of residues allowed per loop: the BC loop was allowed to range between 4 and 10 residues to replace the motif PAVT (residues 25 to 28); the FG loop was allowed to range between 7 and 13 residues to replace the motif GRGDPASSK (residues 77 to 86). The DE loop size is not altered from the “wild-type” (PDB 1TTG).

Next, instead of a free “hallucination”, these generated Mob scaffolds were used as the template for binder design using AlphaFold backpropagation (i.e., AfDesign) implemented in ColabDesign (https://github.com/sokrypton/ColabDesign). The sequence optimization was performed in two stages using the default *design_pssm_semigreedy* protocol, as described previously (Pacesa et al., 2025). Briefly, at each step, a loss based primarily on contacts with the target is calculated using the resulting predicted structure. In the first stage, the Mob sequence is represented as logit inputs or as a position-specific scoring matrix (PSSM) and optimized in a continuous sequence space. The loss is backpropagated through the input sequence, allowing gradients to be computed. Next, in the second stage, the sequence is first converted to a one-hot discrete representation. At each step, 10 random mutations are introduced to the sequence based on the probability distribution from the previous stage, with a bias toward lower pLDDT residues.

No cysteines were allowed during design to ensure easy, high-yield Mob expression in bacteria. The inference loss combined pLDDT, interface contacts, and SolubleMPNN loss with learned weights. Optional losses include a contact loss applied to the desired epitope to drive Mob contacts with the user-defined epitope residues, as well as a sequence identity loss on the Mob framework to reduce amino acid changes in these regions. However, the design pipeline did not apply contact loss when generating designs against RBD. To further expand Mob diversity while improving solubility, SolubleMPNN was applied with a sampling temperature of 0.2 to redesign the interface, defined as residues with Cβ within 7 Å of the target. The first 24 residues of the Mob at the N-terminus remain fixed during all design stages.

All sequences were then ranked using a strategy based on AF2Rank (Roney and Ovchinnikov, 2022). This framework performs AF2-based predictions, with a bias towards a provided template. This pipeline mitigates AF2’s lower predictive performance for this type of interface. To reduce bias when providing a template structure, residues at the interface (within 7 Å of the opposite chain) were masked, and interchain templates were removed. A filtering strategy for selecting the top results followed the following thresholds: pLDDT ≥ 90; ipTM ≥ 0.75; and RMSD between the structure designed with AfDesign and the prediction ≤ 2.5 Å.

### In silico Mob binder design refinement and selection

Starting from the 7 Mobs found in the yeast display screening, we have applied ProteinMPNN mutations (using the soluble weights) in 5 different strategies:

1. mutations outside the interface region (defined as any residue within 7Å of the target protein), excluding the 24 N-terminal residues, with the primary objective of trying to optimize the production yield;
2. mutations only at the interface region to primarily improve binding affinity;
3. mutations only at the interface region (with a maximum of 6 mutations applied to a random subset of residues), to try to refine binding affinity with different mutated residues to promote increased sequence diversity;
4. Combining strategies 1) and 2);
5. Combining strategies 1) and 3).

Across all strategies above, in addition to avoiding mutations to cysteines, we did not mutate existing prolines to avoid promoting significant conformational changes relative to the design starting points. In order to increase sequence diversity, the ProteinMPNN temperature was set at 0.7.

Those sequences were then predicted with AF3 in complex with the target (RBD). The Mobs were predicted in a single-sequence mode. The RBD was predicted with both multiple sequence alignments and structural templates. Results with ipTM ≥ 0.85 and RMSD (target-target alignment between the starting point and the design using PyMOL’s Python package, and then complex-complex (before and after the AF3 prediction) RMSD ≤ 1.5 Å, as well as a PBEE-predicted affinity below 1e-08, were kept (Chaves et al., 2025b). Sequences with fewer than two mutations to any accepted result were removed. The top 50 sequences (ranked by ipTM) were selected.

### Molecular Dynamics simulations

Atomistic MD simulations were performed using the GROMACS 2021.7 (Lindahl et al., 2023) suite and the Amber14sb force field (Maier et al., 2015). The systems were solvated in a dodecahedral box with water, using the transferable intermolecular potential three-point (TIP3P) model (Mark and Nilsson, 2001). The distance between the complex and the box limits was set to a minimum of 2.5 nm. Chloride (Cl^-^) and sodium (Na^+^) ions were added, aiming to reach an ionic strength of 0.15 M while neutralizing the system.

The simulation systems underwent energy minimization protocols, during which we used the steepest-descent method for up to 50,000 steps, during which position restraints were applied to the heteroatom positions, holding them at the specified coordinates with a force constant of 1,000 kJ/mol/nm in the X, Y, and Z directions. Then, the system was initialized through four distinct stages, each lasting 100 picoseconds (ps). In the first stage, all non-hydrogen atoms were restrained with a force constant of 1,000 kJ/mol/nm, and the initial velocities were generated from a Maxwell-Boltzmann distribution. The temperature was controlled using the Berendsen thermostat to keep the simulation systems at 300 K, with a temperature coupling constant of 0.01 ps applied independently to the protein and solvent atoms. No pressure coupling was applied in this step. In the following step, the temperature coupling constant was increased to 0.1 ps. In the third stage, we kept the previously defined temperature-coupling parameters and included pressure coupling via the Berendsen algorithm, with a pressure-coupling constant of 1.0 ps applied isotropically. In the fourth and final stage, we reduced the position restraints from all non-hydrogen atoms to only the Cα atoms, using the same force constant. The temperature algorithm was replaced with V-rescale, with a temperature-coupling constant of 0.1 ps. The pressure algorithm was replaced with the Parrinello-Rahman, with a pressure-coupling constant of 2.0 ps.

Before starting the production part of the simulations, a 100 ns slow-release step was performed. In the 50 ns of this step, we replicated all conditions from the last initialization step. In the second step, we applied restraints only to the Cα atoms of regions with defined secondary structure (as determined by the Python *pydssp* package), using the same force constant of 1,000 kJ/mol/nm.

During the production simulation, the conditions were kept as in the last step of the slow-release stage, but with all restraints removed. During the initialization and production runs, the equations of motion were integrated with a time interval of 2 femtoseconds (fs). Long-range electrostatic interactions were treated through the Particle Mesh Ewald (PME) method, with a grid spacing of 0.12 nm and cubic interpolation. The neighbor list was updated every 20 steps, and the Verlet cutoff scheme used a real-space cutoff radius of 0.8 nm. The LINCS algorithm was leveraged to constrain hydrogen-containing bonds. Three replicates of each system were simulated for a total time of 1 microsecond (μs).

Results from these simulations were analyzed using Python and MDAnalysis (Michaud-Agrawal et al., 2011). The bootstrap of the RMSD values was calculated at 95.45% confidence intervals, assuming a normal distribution, based on bootstrap sampling of the mean across the 3 generated replicates.

### Molecular Mechanics Poisson–Boltzmann Surface Area (MMPBSA)

The MMPBSA analysis was carried out through the gmx_MMPBSA tool (Valdés-Tresanco et al., 2021), employing the single-trajectory approximation. We extracted the frames from the last 500 ns (500 frames saved at 1 ns intervals) of each simulation. Calculations were performed using the Amber14sb force field at 300K. The Poisson-Boltzmann solvation model was used, with the internal dielectric constant set to 1.0, the external dielectric constant set to 80.0, and an ionic strength of 0.150 M. The nonpolar solvation term was calculated based on solvent accessible surface area (SASA). Conformational entropy was not explicitly included in the binding free energy calculations due to significant statistical variability, which can lower the accuracy of binding free energy estimates (Adediwura et al., 2024; Piccardi et al., 2025). Per-residue energy decomposition was conducted on residues within 6 Å of the binding interface to identify key contributors to binding affinity and to better understand interface interactions.

### Yeast surface display library preparation

The DNA oligopool corresponding to the 970 designs was purchased from Twist Bioscience, containing 17-bp overhangs. The oligopool was resuspended in H_2_O to 10 ng μl^−1^ and amplified by two rounds of PCR: (1) for amplification of the library using the 17-bp overhangs (Amp_Fw and Amp_Rev primers, 61 °C annealing for 30 s, 72 °C extension time for 1 min, 15 cycles); and (2) for addition of 45-bp homology with the yeast display vector (Ext_Fw and Ext_Rev primers, 57.5 °C annealing for 30 s, 72 °C extension time for 1 min, 15 cycles). The PCR product was analyzed by agarose gel electrophoresis and subsequently purified, desalted, and co-transformed with a linearized pCTcon2 vector (1:5 vector:insert) into *Saccharomyces cerevisiae* EBY100 cells by electroporation, as previously described (Chao et al., 2006). The pCTcon2 vector contains a fixed N-terminal sequence corresponding to the first 21 residues of the Mob designs, described in PDB 1TTG. The transformation efficiency was at least 4 × 10^6^. Transformed yeast cells were grown for approximately 48 h at 30 °C in SDCAA medium (1.25 g L^−1^ of yeast nitrogen base, 5 g L^−1^ of casamino acids, 13.6 g L^−1^ of Na2HPO4, 8.5 g L^−1^ of NaH2PO4, and 20 g L^−1^ of glucose in 1 L of deionized water) supplemented with 1% of penicillin and streptomycin.

### Yeast surface display

Transformed yeast cells were subcultured once in SDCAA medium prior to overnight induction with SGCAA medium (SDCAA with galactose replacing glucose) at 30 °C. Induced cells were washed with phosphate-buffered saline (PBS) supplemented with 0.1% bovine serum albumin (PBS-B), and incubated with 1 µM of RBD in PBS-B for 2 h at 4 °C. Cells were then washed with PBS-B and labelled with an FITC-conjugated goat anti-HA tag antibody (Bethyl, A190-138F; display tag; 1:100 dilution) and a PE-conjugated goat anti-human Fc antibody (Invitrogen, 12-4317-87; binding tag; 1:100 dilution) for 30 min at 4 °C. Cells were washed with PBS-B, resuspended in an appropriate volume of PBS-B, and sorted on a Cytek Aurora CS in “purity” mode. The sorted yeast cells were grown in SDCAA medium for 48 h, incubated with 500 nM RBD in PBS-B, and labeled for a second sorting round (in “Multiway” mode) to increase selection stringency and enrich for high-affinity binders. Cells obtained after the second and third sorting round were plated on SDCAA agar plates, and 20 individual colonies from different sorting gates were picked for sequencing. A third and final sorting at 50 and 100 nM enabled to find the 7 *in silico* designed Mobs.

### Primers PCR

*Amp_Fw_RBD: 5’ TCCAACTTCTTTGTTGATTAGC 3’*

*Amp_Rev_RBD: 5’ TGGGTATGAACCGGATCC 3’*

*Ext_Fw_RBD: 5’TCCAAGGGATTTGGAAGTTGTTGCTGCTACTCCAACTTCTTTGTTGATTAGC 3’*

*Ext_Rev_RBD: 5’ CTATTAAGCGTAGTCTGGAACGTCGTATGGGTATGAACCGGATCC 3’*

### Recombinant Protein Production

pET28-based plasmids encoding each of the selected Mob designs with an N-terminal hexahistidine (6xHis-) tag were purchased from GeneCust (Boynes, France). The produced protein sequences can be retrieved from **Table S10**. The plasmids were transformed into *Escherichia coli* (*E. coli)* BL21(DE3) by heat shock and bacteria were plated on Luria Broth (LB) agar (GRiSP Research Solutions, Porto, Portugal) plates, supplemented with kanamycin (50 μg/mL) (Carl Roth, Karlsruhe, Germany), incubated overnight at 37 °C. A single colony was selected for overnight growth in 5 mL LB medium (GRiSP Research Solutions), supplemented with kanamycin (50 μg/mL) (LB + Kan), at 37 °C and 160 rpm. Main cultures (100-200 mL) were prepared with 1% *(v/v)* of the overnight grown pre-inoculum in LB Autoinduction medium (GRiSP Research Solutions) supplemented with kanamycin (50 μg/mL) (LB AIM + Kan) and incubated at 37 °C until an optical density at 600 nm (OD600) of 2.0. This was followed by an overnight incubation at 18 °C, 140 rpm. Cell pellets were collected by centrifugation (10,000 ×g for 7 minutes at 4 °C) and stored at −20 °C. Cell pellets were resuspension in lysis buffer (20 mM HEPES, pH 7.5, 300 mM NaCl, 20 mM imidazole, 10% *(v/v)* glycerol, 1 mg/ml lysozyme, 50 U/mL benzonase, 2 mM MgCl₂, 1:500 protease inhibitor cocktail) and cell disruption was achieved by sonication (UP200S sonicator; Hielscher Ultrasonics, Teltow, Germany): 8 cycles, each consisting of 30 seconds of sonication (cycle of 0.5 and amplitude 60) followed by 30 seconds of rest on ice. The resulting suspension was centrifuged at 45,000 ×g at 4 °C for 30 minutes, and the supernatant was collected and filtered through a 0.45 μm syringe filter (VWR, PA, USA).

YSD-discovered Mobs purification was done by affinity HisGravitrap and PD10 desalting columns (Cytiva, MA, USA). Soluble protein fractions were immobilized onto HisGravitrap columns equilibrated with 20 mM HEPES pH 7.5, 300 mM NaCl, 20 mM imidazole, 10% *(v/v)* glycerol (binding buffer, BB). Non-specifically/low affinity bound proteins were removed by an intermediate elution step (BB with 75 mM imidazole), while Mobs were eluted with 3 mL 20 mM HEPES pH 7.5, 300 mM NaCl, 500 mM imidazole, 10% *(v/v)* glycerol (elution buffer, EB). Eluted Mobs were then buffer exchanged to 20 mM HEPES pH 7.5, 150 mM NaCl with PD10 columns.

Purification of refined Mobs followed a two-step protocol carried out in an ÅKTA Go system (Cytiva). First, soluble protein fractions were loaded onto a 5 mL HisTrap HP column (Cytiva) equilibrated with BB buffer, followed by a 10-column volume (CV) wash step (0% EB buffer). Bound proteins were eluted with a 20-CV linear gradient to 50%EB buffer, followed by a -CV step at 50% EB and a -CV gradient to 100% EB buffer. All steps were performed at 3 mL/min, with a fractionation volume of 3 mL during the gradient step. Mob-containing fractions were analyzed by SDS-PAGE, pooled, concentrated in Amicon® Ultra centrifugal filters (3 kDa MWCO), and further purified by size exclusion chromatography on a Hiload 16/600 Superdex 75 pg column (Cytiva), equilibrated and eluted with 20 mM HEPES pH 7.5 and 150 mM NaCl buffer (1 mL/min flow and 1 mL fractions). Protein purity was confirmed by SDS-PAGE, and concentration was determined by measuring absorbance at 280 nm (Abs 280 nm) with the Shimadzu UV-1800 Spectrophotometer (Shimadzu, Kyoto, Japan), using the theoretical molar extinction coefficient at 280 nm and the molecular weight computed with the ExPASy ProtParam tool (https://web.expasy.org/protparam/). When necessary, Mob solutions were filter-sterilized into sterile, low-binding microtubes inside a laminar flow hood using sterile AcroDisc Supor 0.2 µm 13mm filters (Cytiva).

### Screening of 50 refined Mobs

A protocol was developed to produce and characterize Mobs on a small scale for thermal stability and target-binding activity, set up in multi-well plates, ultimately allowing screening of up to 96 designs in less than 2 weeks.

Chemically competent *E. coli* BL21 (DE3) in 96-well plates (ThermoFisher) were transformed with pET-based plasmids encoding the refined Mobs (purchased from Genecust). Heat shock (45 seconds at 42 °C) was performed in an Applied Biosystems 2720 Thermal Cycler, and the plate was subsequently placed on ice. The cell suspensions (20 µL/well) were transferred to deep-well 96-well plates (VWR) containing 500 µL SOC medium (GRISP Research Solutions) and left to recover for 2 h at 37 °C and 200 rpm, then transferred thereafter to 24-well plates (VWR) containing 2 mL LB + Kan and incubated overnight at 37 °C and 200 rpm. The following day, 75 µL of pre-inocula were transferred to 5 mL LB AIM + Kan in new 24-well plates. When the culture reached an OD600nm of 2.0 (monitored in a Fluostar Optima), the incubation temperature was decreased from 37 °C to 18 °C, and the culture was left to grow at 200 rpm for 16 - 18 h. Cells were harvested by centrifugation at 2,200 ×g for 20 min in an Eppendorf 5810R Benchtop centrifuge (Eppendorf, Hamburg, Germany) and stored at −20 °C.

Cells were resuspended in NZYTech Bacterial Cell Lysis buffer (NZYTech, Lisbon, Portugal), the suspensions were transferred to a single deep-well 96-well plate, and incubated at room temperature for 30 min. After centrifugation (2,200 ×g for 30 min, at 4 °C), the cleared supernatants corresponding to the soluble extract were transferred to a MultiTrap 96-well IMAC plate (Cytiva) previously equilibrated with PBS pH 7.4, 150 mM NaCl, 40 mM imidazole. One wash step was performed with PBS pH 7.4, 150 mM NaCl, 75 mM imidazole to elute low-affinity protein binders, and the Mobs were eluted with 130 µL (3 elutions with 43 µL elution buffer, 500 ×g for 2 min at 4 °C between each step). The eluted proteins were then buffer exchanged to PBS pH 7.4 with a MiniTrap (G25) 96-well plate (Cytiva) (eluted by centrifugation (800 ×g for 2 min at 4 °C) and quantified with the BCA method.

The resulting partially purified Mobs were further analyzed for thermal stability and binding activity, as described in the respective sections below: “*Protein thermal stability assessment*” and “*Surface plasmon resonance*”.

### Protein thermal and colloidal stability assessment

Sample monodispersity, thermal denaturation, and aggregation were analyzed in a Prometheus Panta (NanoTemper, Germany). Dynamic light scattering measurements were performed by transferring 10 µL of Mob sample (at ∼1 mg/mL; in triplicate) to Nanotemper standard glass capillaries and performing Size Analysis at 20 °C, with 10 accumulations of 5 seconds each, and 100% laser excitation power. Thermal denaturation was assessed by differential scanning fluorometry (DSF), monitoring the intrinsic tryptophan fluorescence changes at 330 nm and 350 nm as a function of a linear temperature gradient. Thermal aggregation was simultaneously analyzed by light-backscattering (turbidometry, assessing formation of large insoluble aggregates) and dynamic light scattering (DLS, assessing particle size and scattering intensity of soluble aggregates). The capillaries used for Size Analysis measurements were subjected to a linear temperature gradient from 20 to 90 °C at 1 °C/min. Mean values of three replicates are presented. DSF graphs plot fluorescence emission (either the 350 nm/330 nm ratio or the emission at 330 nm) as a function of temperature, and first-derivative graphs were calculated with the manufacturer’s software. T_m_ is defined as the DSF melting temperature, T_on_turbidity_ as the onset of the turbidometry increase in light-backscattering curves, and T_on_scattering_ as the onset of the scattering increase in DLS curves.

### Far-UV circular dichroism spectropolarimetry

Far-UV circular dichroism (CD) spectra of Mobs were acquired in a Jasco-1500 CD spectrometer (Jasco Global, Japan) coupled with a Peltier temperature controller, a six-position sample holder, and a 1-mm optical path quartz cuvette. Measuring parameters were as follows: wavelength range of 260-200 nm, data pitch of 0.1 nm, standard sensitivity, D.I.T. of 2 sec, 2 nm bandwidth, scanning speed of 50 nm/min, 4 accumulations, and measuring temperature of 20 °C. All spectra were offset at 260 nm, and the buffer spectra were subtracted from the Mob spectra.

### Fluidics-free surface plasmon resonance

Binding assays were performed using the fluidics-free surface plasmon resonance (SPR) White FOx equipment (Vixen BIOSYSTEMS). Two types of experiments were carried out: binding to RBD at two fixed RBD concentrations as a screening approach (for the Mobs screening protocol), or affinity constant (K_D_) determination, evaluating Mob binding to several RBD concentrations. For both approaches, Mobs were diluted to 0.5 µg/mL in PBS with 0.01% Tween, pH 7.5 (PBS-T), then immobilized on nickel-coated NTA biosensors for 75 s and then washed to remove excess unbound protein and set the baseline (180 s). For binding screening, Mob-bearing biosensors were transferred to solutions containing 500 or 50 nM RBD (in PBS-T) to monitor association (180 s), and then transferred to PBS-T solutions to monitor dissociation (600 s). For K_D_ determination for selected Mobs, the same association and dissociation times were employed, while RBD binding was assayed for 8 concentrations in the 3.9-500 nM range, from the lowest to the highest, without sensor regeneration steps. Data were fitted locally with single exponential association and decay curves. Association bimolecular rate constants (*k*_on_) were determined by linear regression of observed association rate constants as a function of RBD concentration. Dissociation rates (*k*_off_) were determined by averaging the fitted exponential decay rate constants. Statistical analysis was conducted employing the Jackknife resampling method. All assays were performed at 26 °C.

Competition assays were performed by immobilizing Mobs as described above, and comparing the association and dissociation curves to 100 nM RBD alone with those for a solution containing 100 nM RBD pre-incubated with equimolar ACE2 (in PBS-T).

### Pseudovirus Neutralization Assay

Spike-pseudotyped lentiviral particles were produced by co-transfection of HEK293T cells with the pLEX-GFP reporter plasmid, psPAX2 packaging plasmid, and pCAGGS-Spike WT or variant expression vector, as described previously (Gonçalves et al., 2023). For neutralization assays, Mob samples were serially diluted threefold (nine dilutions) and incubated with pseudoviruses for 1 hour at 37 °C. The virus–Mob mixtures were then added to 293T-ACE2 cells (Alenquer et al., 2021) seeded in 96-well plates at a final multiplicity of infection (MOI) of 0.2. At 48 hours post-infection, GFP fluorescence was quantified using the GloMax Explorer System (Promega). Relative fluorescence units (RFU) were normalized to virus-only control wells (infected cells without Mobs) after subtracting background values from cell-only wells. The IC₅₀ values were determined by fitting neutralization curves with a four-parameter nonlinear regression model (least-squares, unweighted; normalized response) in GraphPad Prism 8.

### Neutralization Assays with Authentic Virus

For neutralization assays with an authentic virus, SARS-CoV-2 B.1 lineage (obtained through BEI Resources, NIAID, NIH: SARS-Related Coronavirus 2, Isolate Germany/BavPat1/2020, NR-52370) was used. SARS-CoV-2 particles were propagated in Vero CCL81 cells (purchased from ATCC, CCL81^TM^) for 3-4 days. The culture medium containing the infectious viral particles was harvested and centrifuged at 450 × g for 15 min to remove cell debris, as described elsewhere (Mendonça et al., 2024). Virus aliquots were stored at −80 °C and titrated by plaque assay using Vero CCL81 cells.

HEK293T-ACE2-30F-PLP2 reporter cells (obtained from the MHRA, thanks to the contribution of Dr. Nicholas Matheson and Dr. Pehuén Pereyra Gerber) were seeded in white 96-well plates at 4 × 10^4^ cells per well the day prior to infection. Mobs were serially diluted 3-fold to 9 dilutions and incubated with SARS-CoV-2 at a final MOI of 0.025 for 1 h at 37 °C before addition to cells. 24 h after infection, firefly (FF) and renilla (RN) luciferase activities were detected using the Dual-Glo Luciferase Assay System (PROME2920, Promega) according to the manufactureŕs instructions. FF/RN values were used to calculate the percentage of infectivity relative to maximum infectivity. The IC_50_ values were calculated using nonlinear regression (inhibitor vs. normalized response, variable slope, least squares, unweighted) in GraphPad Prism 8. Data are represented as means of at least three different independent experiments.

## Supporting information

All extra images/data not included in the main body of the paper.

## ACKNOWLEDGMENTS

The project leading to these results has received funding from “la Caixa” Foundation and FCT – Fundação para a Ciência e a Tecnologia, I.P. under the project code LCF/PR/HP22/52320025 - HR22-00722 “The BioPlaTTAR Platform for the Tailored and Rapid Development of Antiviral Biopharmaceuticals”; by FCT, I.P., through MOSTMICRO-ITQB R&D Unit (10.54499/UID/04612/2025, UID/PRR/4612/2025), Research Unit iNOVA4Health (UID/04462/2025), LS4FUTURE Associated Laboratory (10.54499/LA/P/0087/2020), GIMM Research Unit (doi.org/10.54499/UID/06357/2025), GIMM Associate Laboratory (LA/P/0082/2020) [doi.org/10.54499/LA/P/0082/2020], and grants 10.54499/2021.09343.BD; 10.54499/2023.08460.CPCA.A1 and 2025.09577.CPCA.A3; by EvaMobs, a HORIZON Europe Framework Programme, under reference HORIZON-HLTH-2023-DISEASE-03-04, GA 101137419; by the Recovery and Resilience Plan (UID/PRR/04462/2025 and UID/PRR2/04462/2025), funded by the European Union – NextGenerationEU; and by an EMBO Scientific Exchange Grant (ID 10333). Additionally, this work was supported by GIMM-CARE (funded by the European Union under grant agreement No. 101060102. GIMM-CARE is co-funded by the Portuguese Government, FCT, ARICA – Investimentos, Participações e Gestão, Jerónimo Martins, the Gulbenkian Institute for Molecular Medicine, and CAML – Lisbon Academic Medical Centre) [doi.org/10.3030/101060102].

Recombinant soluble human ACE2 was a kind gift of Rute Castro and Paula Alves from iBET.

The SARS-CoV-2 B.1 lineage (obtained through BEI Resources, NIAID, NIH: SARS-Related Coronavirus 2, Isolate Germany/BavPat1/2020, NR-52370) was obtained from the MHRA, thanks to the contribution of Dr Nicholas Matheson and Dr. Pehuén Pereyra Ger.

## CODE AVAILABILITY

The MobGen code will be made freely available upon final publication.

