## Supplementary material for "De novo design of antiviral monobody binders": All extra images/data not included in the main body of the paper.

This section will contain all the data regarding protein production and biophysical characterization, binding affinity and neutralization activity assays described throughout the work, as well as any MD analysis not presented in the main body of text.

**Table S1:** Production and characterization of YSD-derived RBD-binding monobodies (Mobs) before (green) and after (blue) addition of stabilizing C-terminal tail. Production yields were calculated as mg of pure protein (after the final size-exclusion chromatography step and the following filter sterilization) divided by the culture volume. Thermal denaturation and aggregation were analyzed with a Nanotemper Prometheus Panta, thereby estimating the melting temperature ($T_m$) by DSF, and the onset of aggregation temperature for large insoluble aggregates ($T_{on\_tubidity}$, obtained from light backscattering turbidometry) and/or soluble aggregates ($T_{on\_scattering}$ , obtained by DLS).

| Mob ID | Yield (mg/L culture) | $T_m$ (°C) | $T_{on\_turbidity}$ (°C) | $T_{on\_scattering}$ (°C) |
|---|---|---|---|---|
| **2515** | 63 | 52.5 ± 0.1 | no | 47.2 ± 1.0 |
| **2550** | 6 | 52.2 ± 0.2 | no | n.d. |
| **2563** | 14 | 42.8 ± 0.4 | no | 32.3 ± 1.3 |
| **2604** | 6 | n.d. | no | n.d. |
| **2683** | 36 | 66.2 ± 0.6 | no | n.d. |
| **2689** | 30 | 52.0 ± 0.1 | no | 50.4 ± 0.6 |
| **2950** | 10 | 45.4 ± 0.1 | 61.4 ± 0.5 | n.d. |
| **2515T** | 164 | 59.0 ± 0.1 | no | 53.0 ± 1.0 |
| **2550T** | 65 | 55.7 ± 0.1 | no | 50.3 ± 0.3 |
| **2563T** | 86 | 61.1 ± 0.1 | no | 54.4 ± 0.9 |
| **2604T** | 55 | 38.3 ± 0.1 | no | 30.6 ± 0.6 |
| **2683T** | 250 | 68.3 ± 0.1 | no | 63.7 ± 0.9 |
| **2689T** | 156 | 56.3 ± 0.1 | no | 51.5 ± 0.2 |
| **2950T** | 114 | 46.7 ± 0.7 | 39.8 ± 3.5 | 38.4 ± 1.8 |

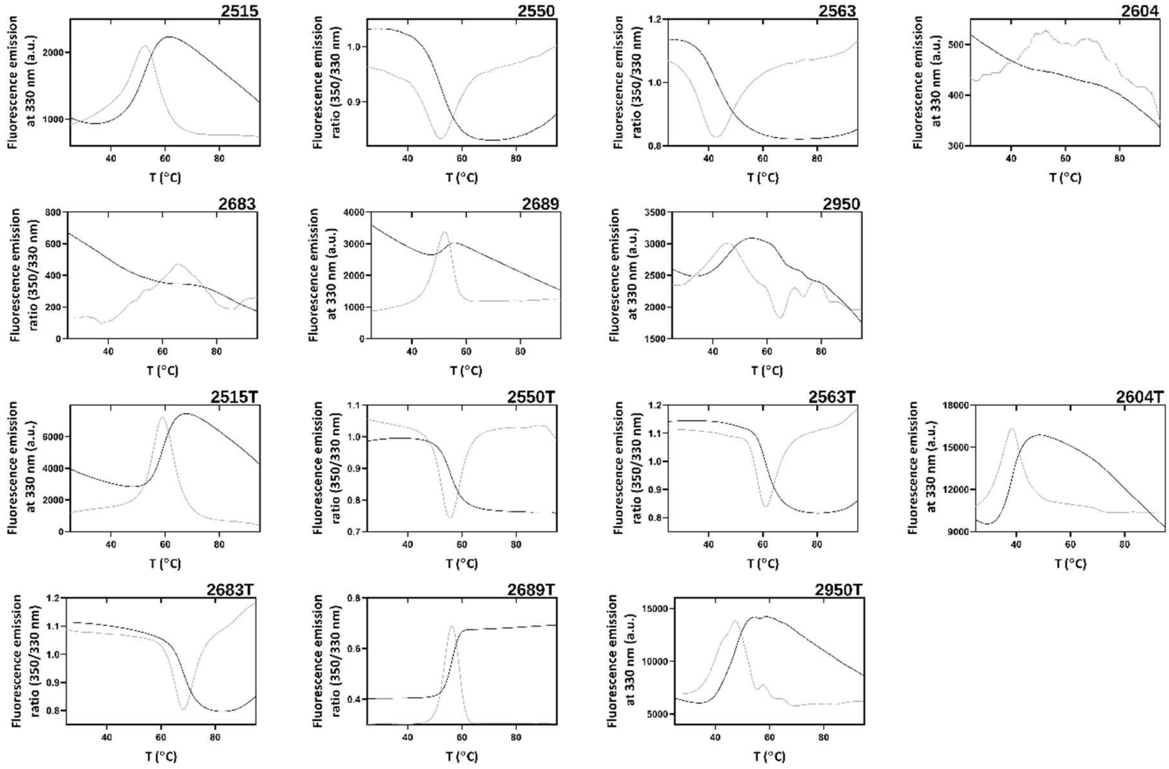


**Figure S1:** Thermal denaturation profiles for YSD-discovered Mobs before and after addition of stabilizing C-terminal tail, assessed by differential scanning fluorometry. Black line, fluorescence emission (ratio between fluorescence emission at 350 and 330 nm or single emission at 330 nm); grey line, first derivative; dashed line marks the melting temperature ($T_m$) value for each Mob.

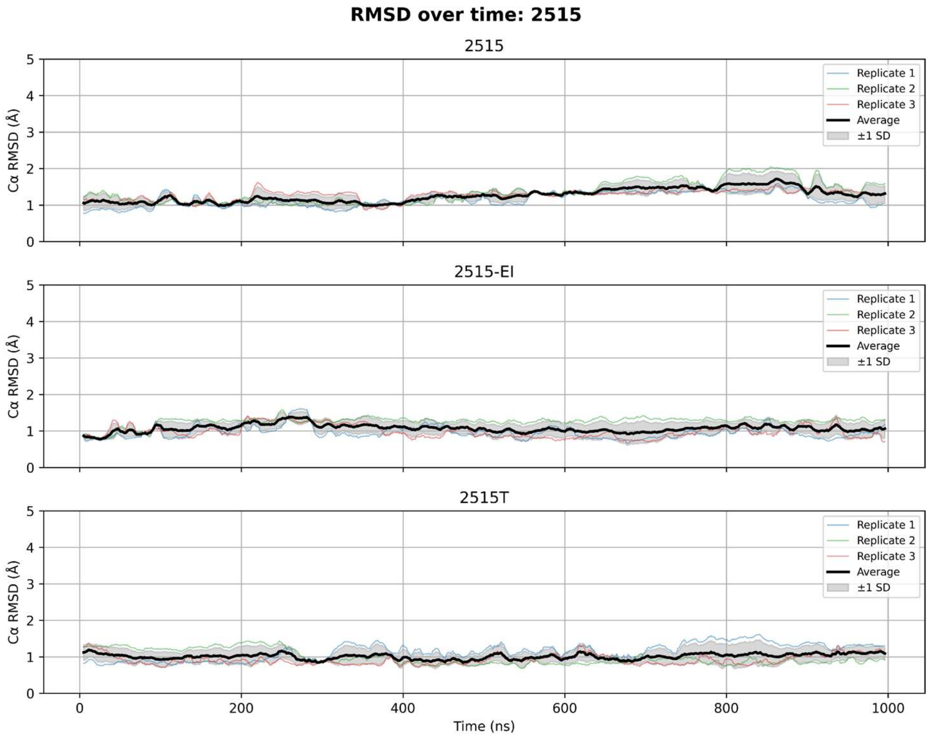


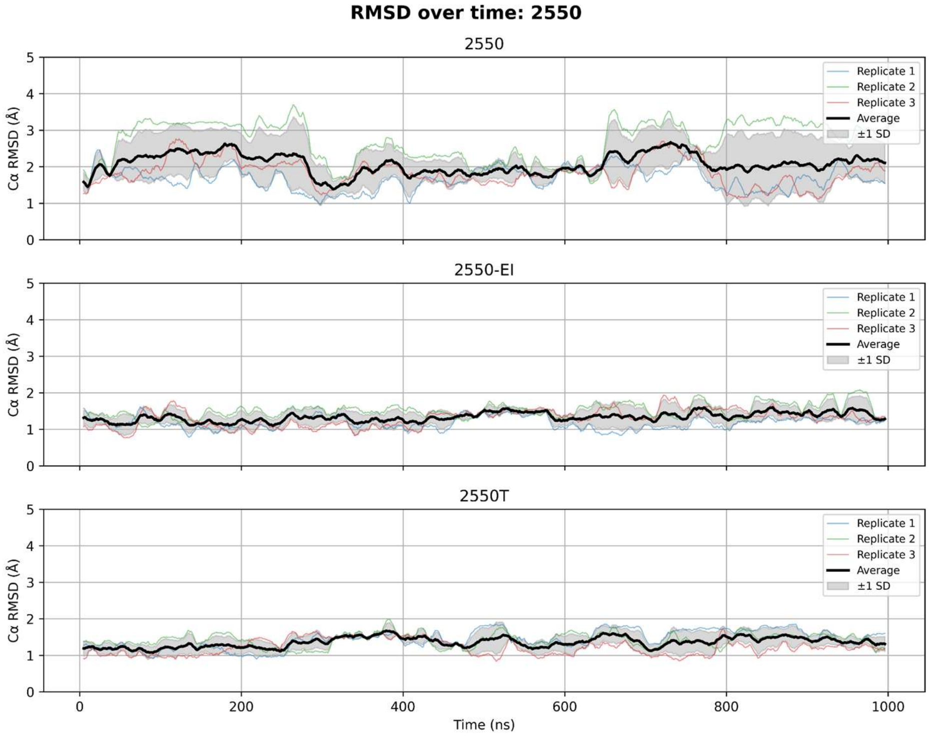


**Figure S2:** RMSD of two YSD-discovered Mobs - 2515 and 2550 - throughout the MD simulation time (1μs), across 3 replicates. These Mobs were simulated without the C-terminal putatively stabilizing tail, with the truncated version of the C-terminal tail ("EI"), and with the full tail ("T"). To compare between simulation conditions, the RMSD is calculated with all residues until the end of the G β-strand (excluding the tails).

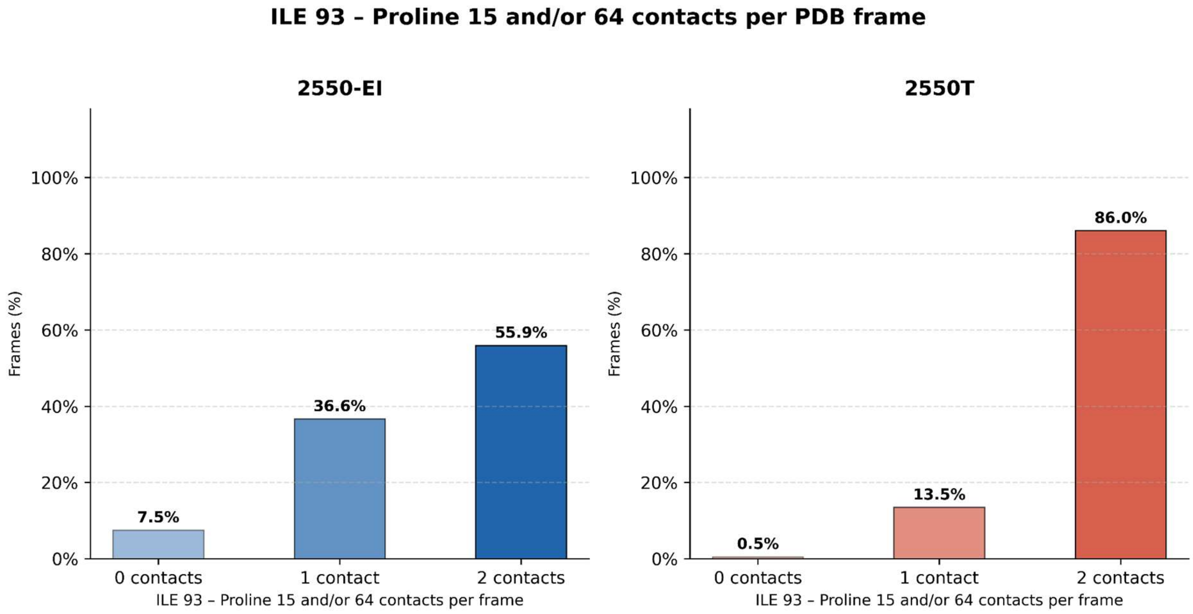


**Figure S3:** Percentage of MD frames in which contacts between isoleucine 93 of Mob-2550-EI and Mob-2550T and proline residues at positions 15 and 64 are observed. 1,000 frames were extracted from each of the 3 simulation replicates (one every 1 ns), for a total of 3,000 frames per Mob. Contacts were identified using MDAnalysis (Michaud-Agrawal et al., 2011), and defined as any heavy-atom distance ≤ 4 Å between the isoleucine and either proline residue.

**Table S2:** Middle-throughput screening of ProteinMPNN-refined Mobs. Protein purification yield was estimated by quantitating the total protein in mg (after the one-step 96-well plate affinity chromatography and buffer exchange column) and dividing by the culture volume. Thermal denaturation analyzed with a Nanotemper Prometheus Panta allowed estimating melting temperature ($T_m$) values. RBD binding was screened by SPR on a Vixen Biosystems White FOx, by immobilizing the Mobs on Ni-NTA sensors, and testing their binding to 50 nM or 500 nM RBD. The Binding Signal at the lowest concentration (50 nM) is herein categorized into three types, based on the signal presented in **Figure S4**: "No / Low" signal; "Medium" signal; or "Strong" signal. Based on all three criteria, selected Mobs are marked in Bold.

| Mob ID | Yield (mg/L culture) | $T_m$ (°C) | SPR Binding Signal at lowest concentration (50 nM) |
|---|---|---|---|
| 9003T | 48 | 49.7 ± 0.3 | No / Low |
| 9004T | 122 | 43.6 ± 0.1 | No / Low |
| 9005T | 23 | 44.0 ± 0.2 | No / Low |
| 9006T | 55 | 72.3 ± 0.1 | No / Low |
| **9007T** | **60** | **74.1 ± 0.1** | **Strong** |
| 9008T | 72 | 52.8 ± 0.1 | No / Low |
| 9009T | 54 | 52.1 ± 0.1 | Medium |
| 9010T | 79 | 51.6 ± 0.1 | No / Low |
| **9011T** | **99** | **54.7 ± 0.3** | **Strong** |
| 9012T | 231 | 68.7 ± 0.1 | No / Low |
| **9013T** | **54** | **65.8 ± 0.1** | **Strong** |
| 9014T | 60 | 53.1 ± 0.1 | No / Low |
| 9015T | 18 | 67.5 ± 0.1 | No / Low |
| **9016T** | **51** | **61.6 ± 0.1** | **Strong** |
| 9017T | 26 | 64.2 ± 0.1 | No / Low |
| **9018T** | **35** | **51.4 ± 0.1** | **Strong** |
| **9019T** | **30** | **67.4 ± 0.1** | **Strong** |
| 9020T | 15 | 58.6 ± 0.1 | Medium |

| | | | |
|---|---|---|---|
| **9021T** | **35** | **50.2 ± 0.1** | **Strong** |
| **9022T** | **34** | **56.3 ± 0.2** | **Strong** |
| 9023T | 17 | 66.6 ± 0.1 | No / Low |
| 9024T | 35 | 66.5 ± 0.1 | Medium |
| **9025T** | **43** | **61.0 ± 0.1** | **Strong** |
| 9026T | 40 | 78.4 ± 0.1 | No / Low |
| 9027T | 36 | 59.6 ± 2.9 | No / Low |
| 9028T | 51 | 56.5 ± 1.5 | No / Low |
| 9029T | 41 | 55.5 ± 0.1 | Medium |
| **9030T** | **25** | **82.3 ± 0.1** | **Strong** |
| 9031T | 47 | 64.8 ± 0.1 | No / Low |
| 9032T | 58 | 50.1 ± 0.1 | No / Low |
| 9033T | 31 | 86.2 ± 0.1 | No / Low |
| 9034T | 52 | 48.8 ± 0.1 | Medium |
| **9035T** | **48** | **61.6 ± 0.1** | **Strong** |
| 9036T | 36 | 59.3 ± 0.1 | No / Low |
| 9037T | 68 | n.d. | No / Low |
| 9038T | 106 | 52.2 ± 0.1 | No / Low |
| 9039T | 40 | 47.3 ± 0.1 | Medium |
| 9040T | 51 | 67.2 ± 0.1 | Medium |
| 9041T | 117 | 53.5 ± 0.1 | No / Low |
| 9042T | 21 | 43.3 ± 0.4 | No / Low |
| 9043T | 57 | 42.5 ± 0.1 | No / Low |
| **9044T** | **3** | **>95** | **Strong** |
| 9045T | 27 | 76.9 ± 0.1 | No / Low |
| 9046T | 77 | 67.4 ± 0.1 | No / Low |

| | | | |
|---|---|---|---|
| 9047T | 62 | 35.9 ± 0.2 | No / Low |
| 9048T | 3 | n.d. | No / Low |
| 9049T | 35 | 59.6 ± 0.1 | No / Low |
| 9050T | 11 | n.d. | No / Low |
| 9051T | 25 | 75.5 ± 0.1 | Medium |
| 9052T | 53 | 58.7 ± 0.1 | Medium |

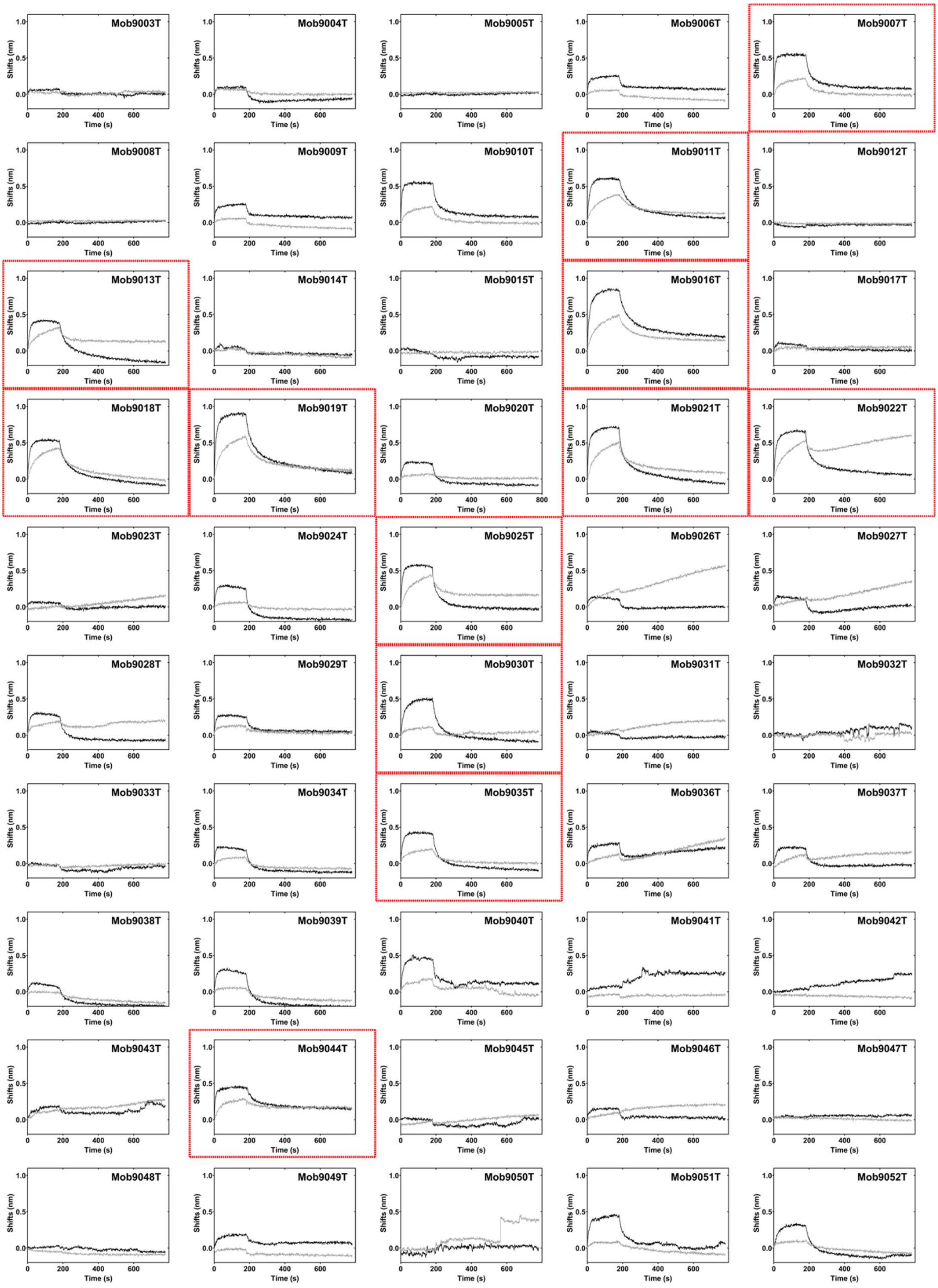
Mob9003T
Mob9004T
Mob9005T
Mob9006T
Mob9007T
Mob9008T
Mob9009T
Mob9010T
Mob9011T
Mob9012T
Mob9013T
Mob9014T
Mob9015T
Mob9016T
Mob9017T
Mob9018T
Mob9019T
Mob9020T
Mob9021T
Mob9022T
Mob9023T
Mob9024T
Mob9025T
Mob9026T
Mob9027T
Mob9028T
Mob9029T
Mob9030T
Mob9031T
Mob9032T
Mob9033T
Mob9034T
Mob9035T
Mob9036T
Mob9037T
Mob9038T
Mob9039T
Mob9040T
Mob9041T
Mob9042T
Mob9043T
Mob9044T
Mob9045T
Mob9046T
Mob9047T
Mob9048T
Mob9049T
Mob9050T
Mob9051T
Mob9052T
Shifts (nm)
Time (s)

**Figure S4:** SPR binding signal from the middle-throughput screening of the 50 ProteinMPNN-refined Mobs, at two different concentrations: black lines, 500 nM; grey lines, 50nM; red box, selected Mobs.

**Table S3**: Production and characterization of ProteinMPNN-refined RBD-binding Mobs. Production yields were calculated as mg of pure protein (after the final size-exclusion chromatography step and the following filter sterilization) divided by the culture volume. Thermal denaturation and aggregation were analyzed with a Nanotemper Prometheus Panta, thereby estimating the melting temperature ($T_m$) by DSF, and the onset of aggregation temperature for large insoluble aggregates ($T_{on\_turbidity}$, obtained from light backscattering turbidometry) and/or soluble aggregates ($T_{on\_scattering}$ , obtained by DLS).

| Mob ID | Yield (mg/L culture) | $T_m$ (°C) | $T_{on\_turbidity}$ (°C) | $T_{on\_scattering}$ (°C) |
|---|---|---|---|---|
| **9007T** | 188 | 73.7 ± 0.1 | no | 67.3 ± 0.1 |
| **9011T** | 212 | 54.4 ± 0.9 | no | 45.7 ± 2.0 |
| **9013T** | 157 | 67.0 ± 0.1 | no | 61.1 ± 0.4 |
| **9016T** | 140 | 63.3 ± 0.1 | no | 57.6 ± 1.1 |
| **9018T** | 105 | 53.6 ± 0.1 | 59.4 ± 0.1 | 39.0 ± 5.2 |
| **9019T** | 165 | 68.7 ± 0.1 | no | 64.6 ± 3.2 |
| **9021T** | 195 | 49.9 ± 0.1 | no | 42.2 ± 0.4 |
| **9022T** | 185 | 57.8 ± 0.1 | no | 46.1 ± 1.3 |
| **9025T** | 240 | 66.4 ± 0.1 | no | 60.0 ± 2.2 |
| **9030T** | 142 | 80.8 ± 0.1 | no | 78.0 ± 1.5 |
| **9035T** | 155 | 62.6 ± 0.1 | no | 58.8 ± 1.3 |
| **9044T** | 168 | 80.8 ± 0.1 | no | 78.4 ± 1.1 |

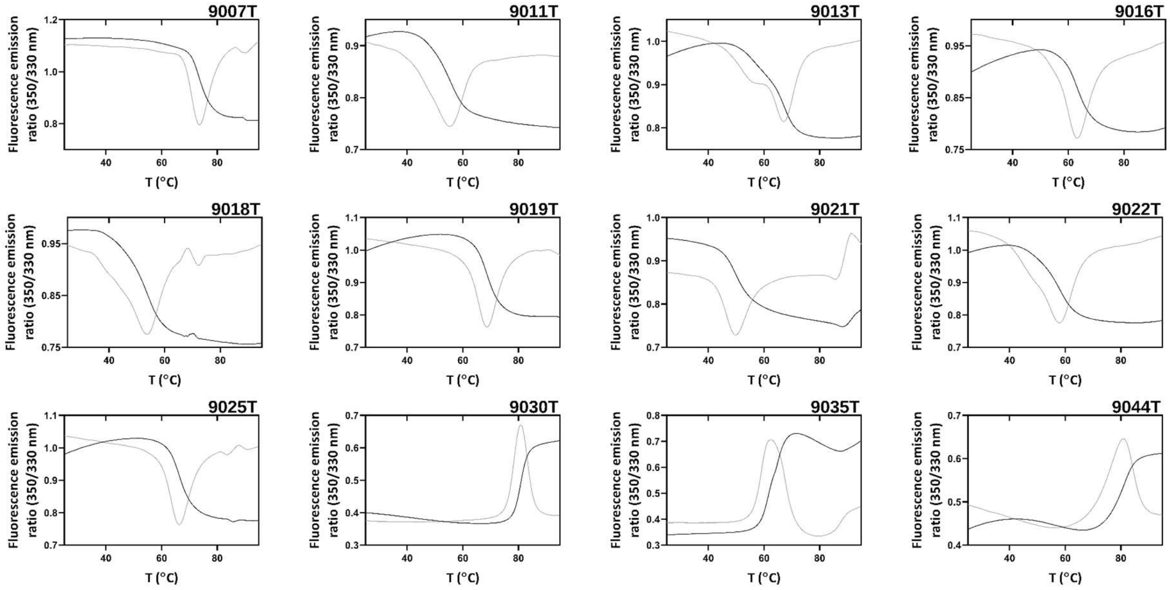


**Figure S5:** Thermal denaturation profiles of the ProteinMPNN-refined Mobs with added tails, assessed by differential scanning fluorometry. Black line, fluorescence emission (ratio between fluorescence emission at 350 and 330 nm or single emission at 330 nm); grey line, first derivative; dashed line marks the melting temperature ($T_m$) value for each Mob.

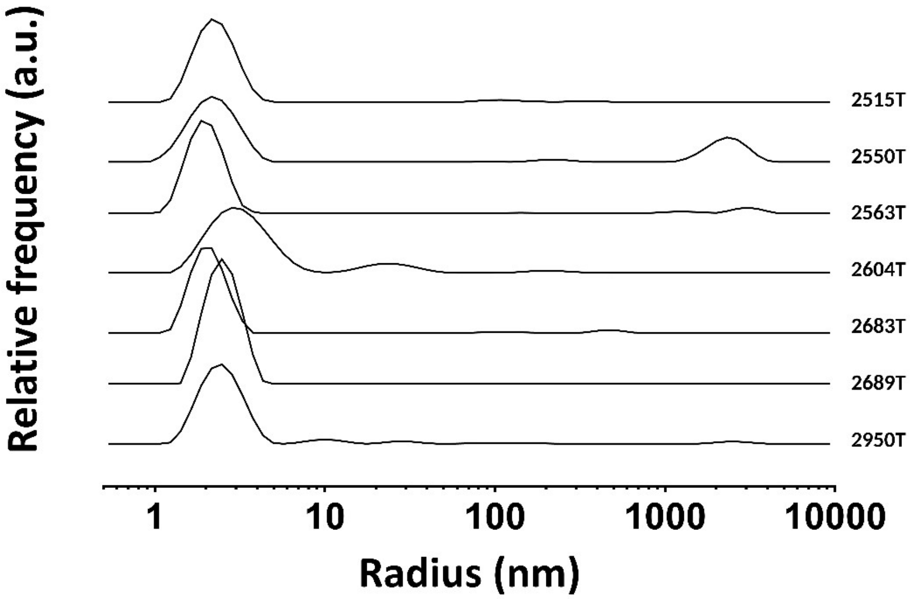


**Figure S6:** DLS profiles of YSD-discovered Mobs after addition of stabilizing C-terminal tail. Population size distribution analysis allows identifying a main population with a hydrodynamic radius around three nm. The majority of the Mobs present a single population profile.

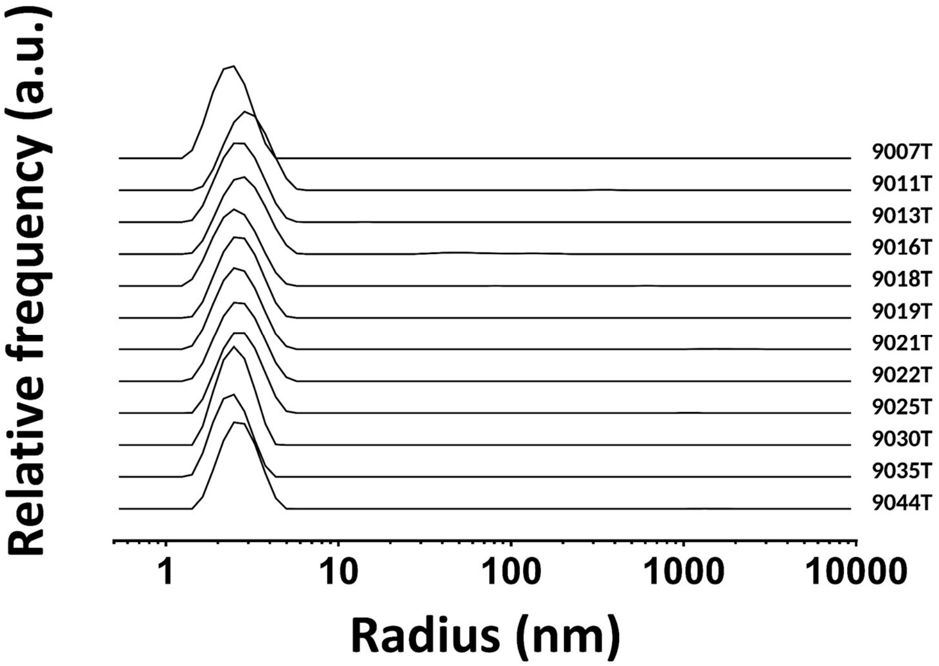


**Figure S7**: DLS profiles of ProteinMPNN-refined Mobs containing the stabilizing C-terminal tail. Population size distribution analysis allows identifying a main population with a hydrodynamic radius around three nm. The majority of the Mobs present a single population profile.

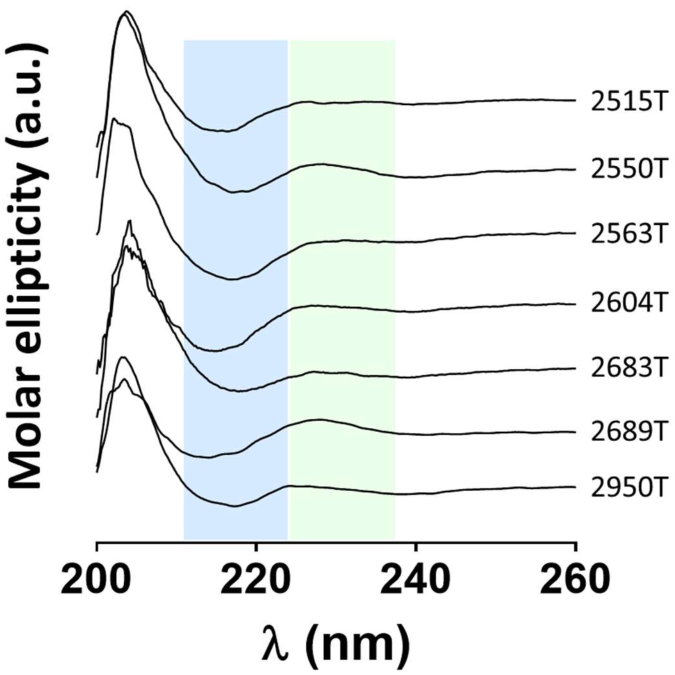


**Figure S8:** Far-UV circular dichroism spectra of YSD-discovered Mobs, after addition of the stabilizing C-terminal tail, confirm predominant β-sheet content and highlight a positive band at 225-230 nm consistent with Mobs CD data. Spectra of Mobs at 0.2 mg/mL in 20 mM HEPES, 150 mM NaCl, pH 7.5 result from four accumulations at 50 nm/min, with a data pitch 0.1 nm, data integration time 2 s, bandwidth 2 nm and $N_2$ flow of 8 L/min. Spectra were offset at 260 nm and the buffer spectrum was subtracted. Stacked spectra are presented for clarity. The blue-shaded box highlights a negative band centered at ~218 nm consistent with β-sheets, while the green-shaded box highlights the 225-230 nm region where an unusual positive band appears in Mobs spectra.

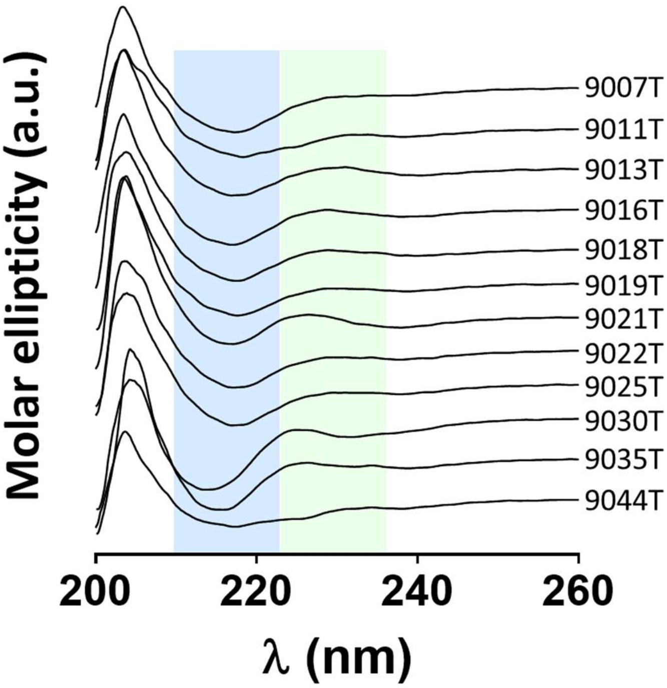


**Figure S9:** Far-UV circular dichroism spectra of ProteinMPNN-refined Mobs, containing the stabilizing C-terminal tail, confirm predominant β-sheet content and highlight a positive band at 225-230 nm consistent with Mobs CD data. Spectra of Mobs at 0.2 mg/mL in 20 mM HEPES, 150 mM NaCl, pH 7.5 result from four accumulations at 50 nm/min, with a data pitch 0.1 nm, data integration time 2 s, bandwidth 2 nm and $N_2$ flow of 8 L/min. Spectra were offset at 260 nm and the buffer spectrum was subtracted. Stacked spectra are presented for clarity. The blue-shaded box highlights a negative band centered at ~218 nm consistent with β-sheets, while the green-shaded box highlights the 225-230 nm region where an unusual positive band appears in Mobs spectra.

**Table S4:** Affinity of refined Mobs (with the added solubility tail and ProteinMPNN-refined) evaluated by SPR, enabling determination of the binding kinetic parameters ($k_{on}$, $k_{off}$, $K_D$).

| **Mob ID** | **$k_{on}$ (±SE) ($M^{-1}·s^{-1}$)** | **$k_{off}$ (±SE) ($s^{-1}$)** | **$K_D$ (±SE) (nM)** |
|---|---|---|---|
| **2515T** | 9.96 ± 6.71 E04 | 6.67 ± 2.02 E-03 | 66.9 ± 49.4 |
| **2550T** | 6.35 ± 0.80 E04 | 9.86 ± 1.67 E-03 | 155.3 ± 32.8 |
| **2563T** | 1.20 ± 0.51 E05 | 2.47 ± 0.54 E-02 | 206.3 ± 98.4 |
| **2683T** | 2.51 ± 0.71 E05 | 1.45 ± 0.33 E-02 | 57.8 ± 20.9 |
| **9007T** | 3.49 ± 0.48 E05 | 7.76 ± 1.20 E-03 | 22.2 ± 4.6 |
| **9011T** | 8.02 ± 0.40 E04 | 8.78 ± 0.97 E-03 | 109.5 ± 13.3 |
| **9013T** | 8.01 ± 4.98 E04 | 1.18 ± 0.51 E-02 | 147.0 ± 111.0 |
| **9016T** | 9.81 ± 2.52 E04 | 9.91 ± 1.04 E-03 | 101.1 ± 28.0 |
| **9018T** | 9.97 ± 0.92 E04 | 7.58 ± 0.93 E-03 | 76.0 ± 11.6 |
| **9019T** | 2.71 ± 1.28 E05 | 8.57 ± 1.14 E-03 | 31.6 ± 15.5 |
| **9021T** | 1.70 ± 0.77 E05 | 9.10 ± 3.49 E-03 | 53.6 ± 31.9 |
| **9022T** | 9.15 ± 3.60 E04 | 6.03 ± 1.37 E-03 | 65.9 ± 30.0 |
| **9025T** | 1.49 ± 0.65 E04 | 6.33 ± 1.26 E-03 | 424.8 ± 203.8 |
| **9030T** | 7.59 ± 1.99 E04 | 7.73 ± 1.57 E-03 | 101.9 ± 33.8 |
| **9044T** | 4.94 ± 2.87 E05 | 1.09 ± 0.24 E-02 | 22.1 ± 13.7 |

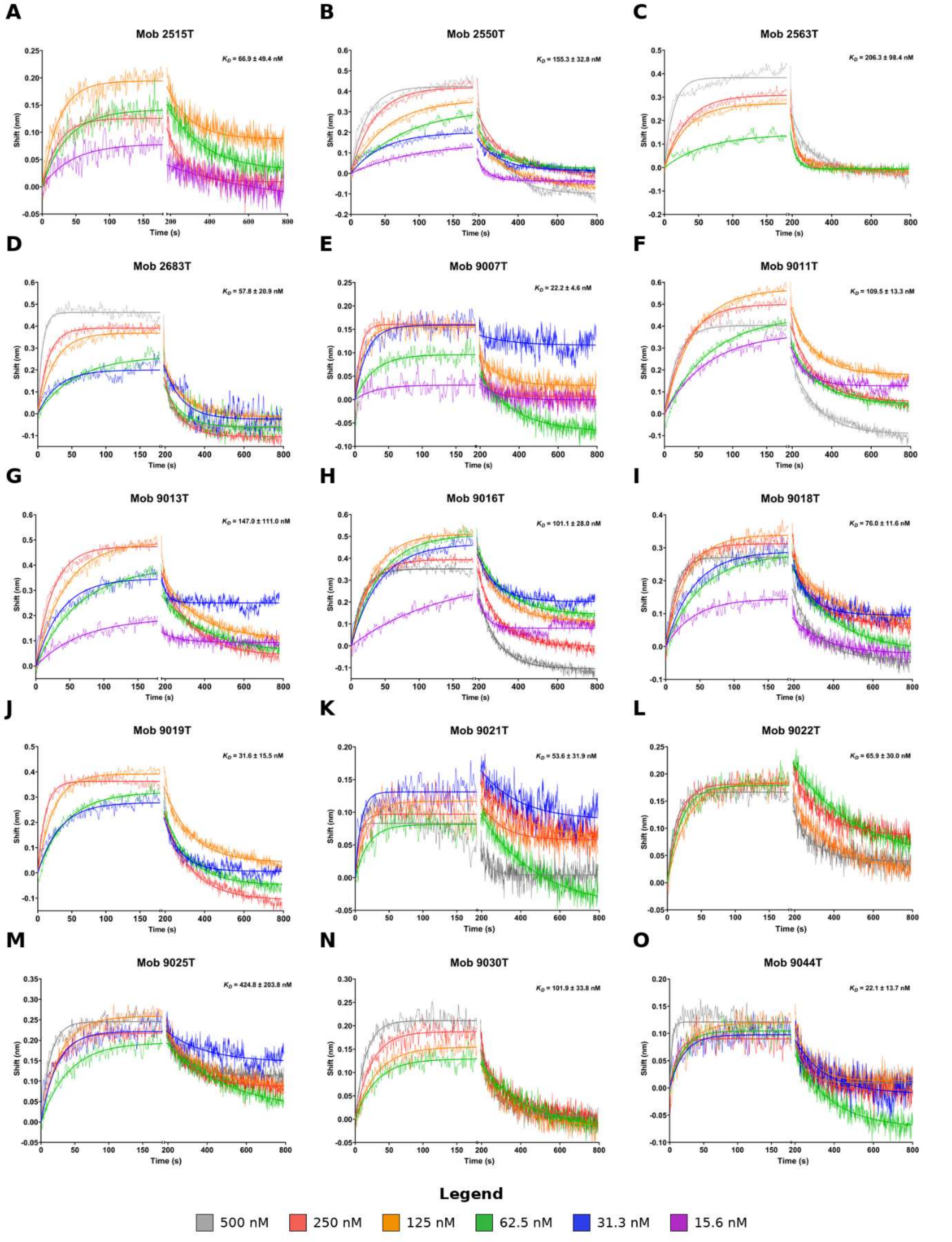


**Figure S10:** SPR sensorgrams for each Mob with measurable $K_D$ values. Association and dissociation curves were measured at different RBD concentrations for each tested Mob.

**Table S5:** Neutralization activity of YSD-derived Mobs against Spike pseudotyped lentiviral particles. Threefold dilution series of each Mob were incubated with the pseudoviruses (MOI 0.2) before being added to 293T-ACE2 cells. GFP fluorescence was measured 48h after infection $IC_{50}$ values are represented as the mean ± Standard Error of the Mean (SEM) from 6 replicates pooled from two independent experiments.

| Mob ID | $IC_{50}$ (±SEM) (μM) (pseudovirus) |
|---|---|
| **2515** | 7.27 ± 0.54 |
| **2550** | >2.33 |
| **2563** | >12.74 |
| **2604** | >1.88 |
| **2683** | >12.54 |
| **2689** | >20.34 |
| **2950** | >6.75 |

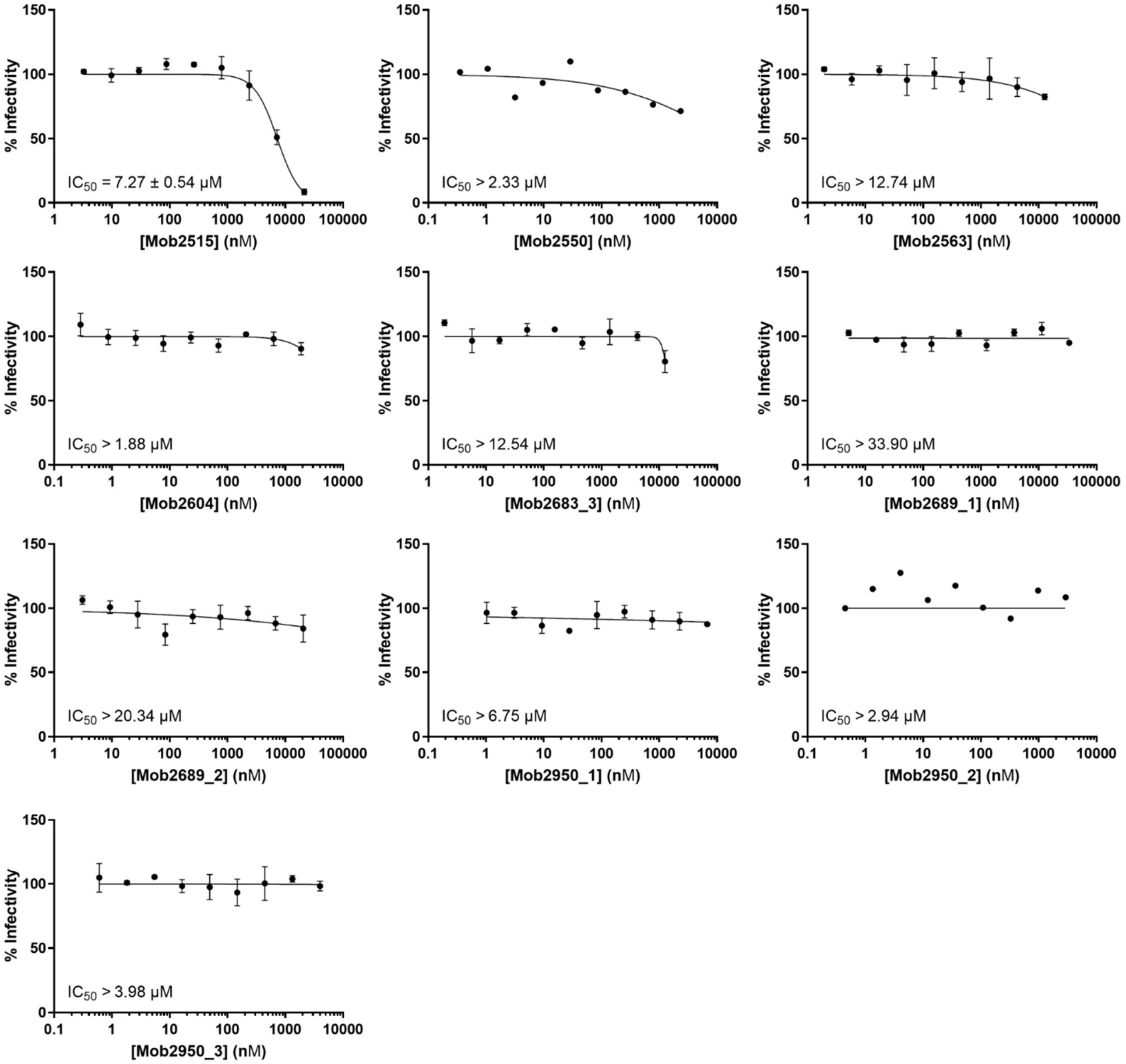


**Figure S11:** One Mobgen-designed Mob (2515) can neutralize the SARS-CoV-2 spike pseudovirus. A dilution series of the indicated Mobs was incubated with Spike pseudotyped virus at an MOI of 0.2 for 1 hour at 37 degrees Celsius, then used to inoculate 293T-ACE2 cells in a 96-well plate. Two days later, fluorescence intensity was measured and infectivity was defined as the percentage of fluorescence relative to cells infected in the absence of Mobs. $IC_{50}$ values were calculated using GraphPad Prism. Data points represent the mean ± SEM of 2 independent experiments, each with 3 technical replicates.

**Table S6:** Neutralization activity of refined Mobs (with the added solubility tail and ProteinMPNN-refined) against Spike pseudotyped lentiviral particles and authentic SARS-CoV-2 B.1 lineage virus. $IC_{50}$ values are represented as the mean ± Standard Error of the Mean (SEM) from 9 replicates pooled from three independent experiments.

| Mob ID | $IC_{50}$ (±SEM) (μM) (pseudovirus) | $IC_{50}$ (±SEM) (μM) (virus) |
|---|---|---|
| **2515T** | 24.42 ± 3.73 | 32.9 ± 4.0 |
| **2550T** | 1.49 ± 0.15 | 3.2 ± 0.4 |
| **2563T** | 4.84 ± 0.42 | 4.7 ± 0.6 |
| **2604T** | 8.21 ± 0.42 | > 40 |
| **2683T** | 62.18 ± 12.57 | > 40 |
| **2689T** | >53.65 | > 40 |
| **2950T** | >37.75 | > 40 |
| **9007T** | 18.54 ± 3.49 | > 40 |
| **9011T** | 2.08 ± 0.28 | 8.8 ± 1.0 |
| **9013T** | 5.71 ± 0.77 | 19.8 ± 2.2 |
| **9016T** | 2.40 ± 0.48 | 8.5 ± 1.4 |
| **9018T** | 17.30 ± 8.65 | 10.0 ± 0.9 |
| **9019T** | 4.99 ± 0.74 | 21.4 ± 2.5 |
| **9021T** | 13.51 ± 3.69 | 40 ± 2.8 |
| **9022T** | 7.82 ± 0.95 | 39.5 ± 3.3 |
| **9025T** | 8.66 ± 0.99 | 35.9 ± 4.3 |
| **9030T** | 19.69 ± 3.89 | > 40 |
| **9035T** | 10.19 ± 1.65 | 27.4 ± 4.1 |
| **9044T** | 19.87 ± 3.13 | > 40 |

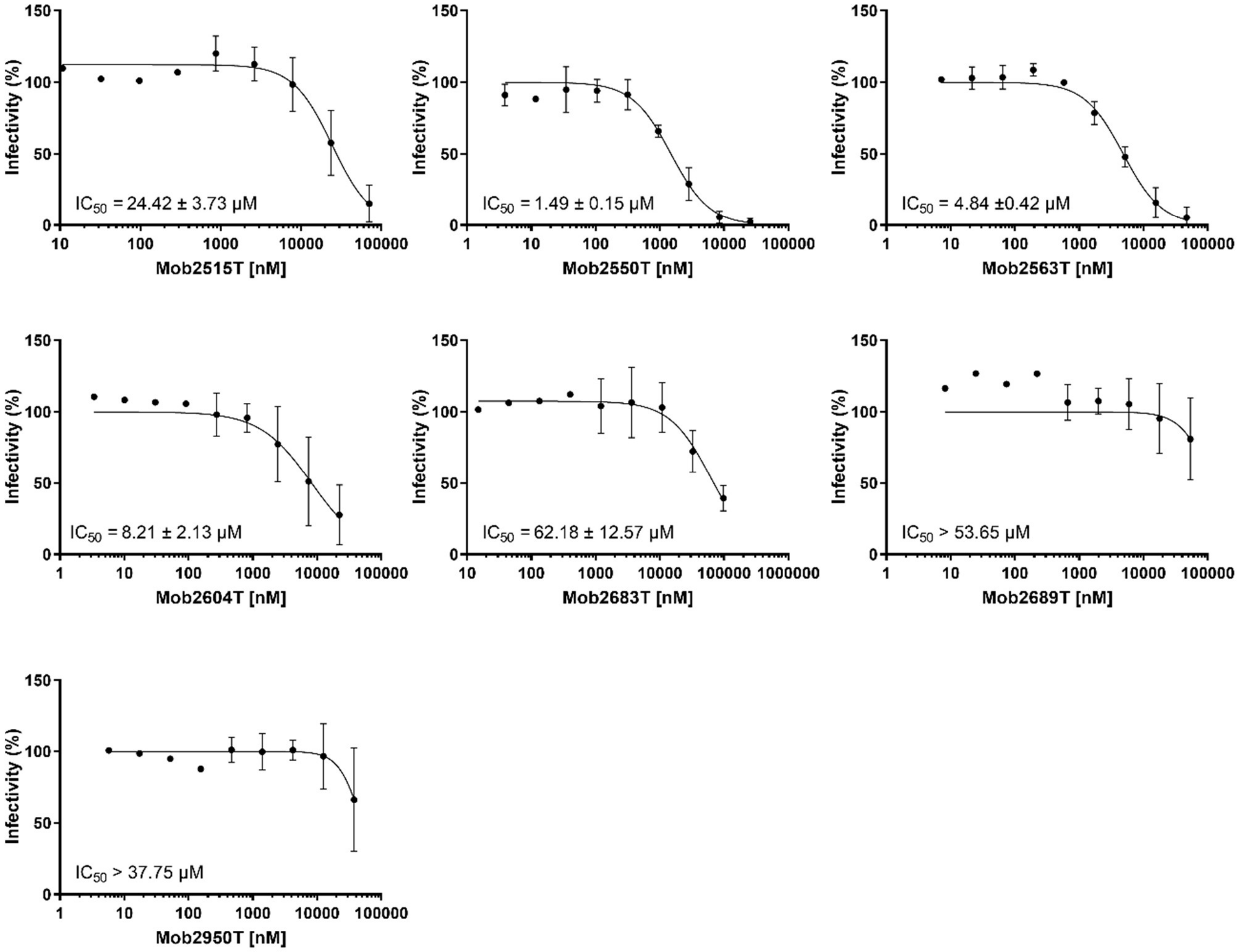

Infectivity (%)
IC50 = 24.42 ± 3.73 µM
Mob2515T [nM]
IC50 = 1.49 ± 0.15 µM
Mob2550T [nM]
IC50 = 4.84 ±0.42 µM
Mob2563T [nM]
IC50 = 8.21 ± 2.13 µM
Mob2604T [nM]
IC50 = 62.18 ± 12.57 µM
Mob2683T [nM]
IC50 > 53.65 µM
Mob2689T [nM]
IC50 > 37.75 µM
Mob2950T [nM]

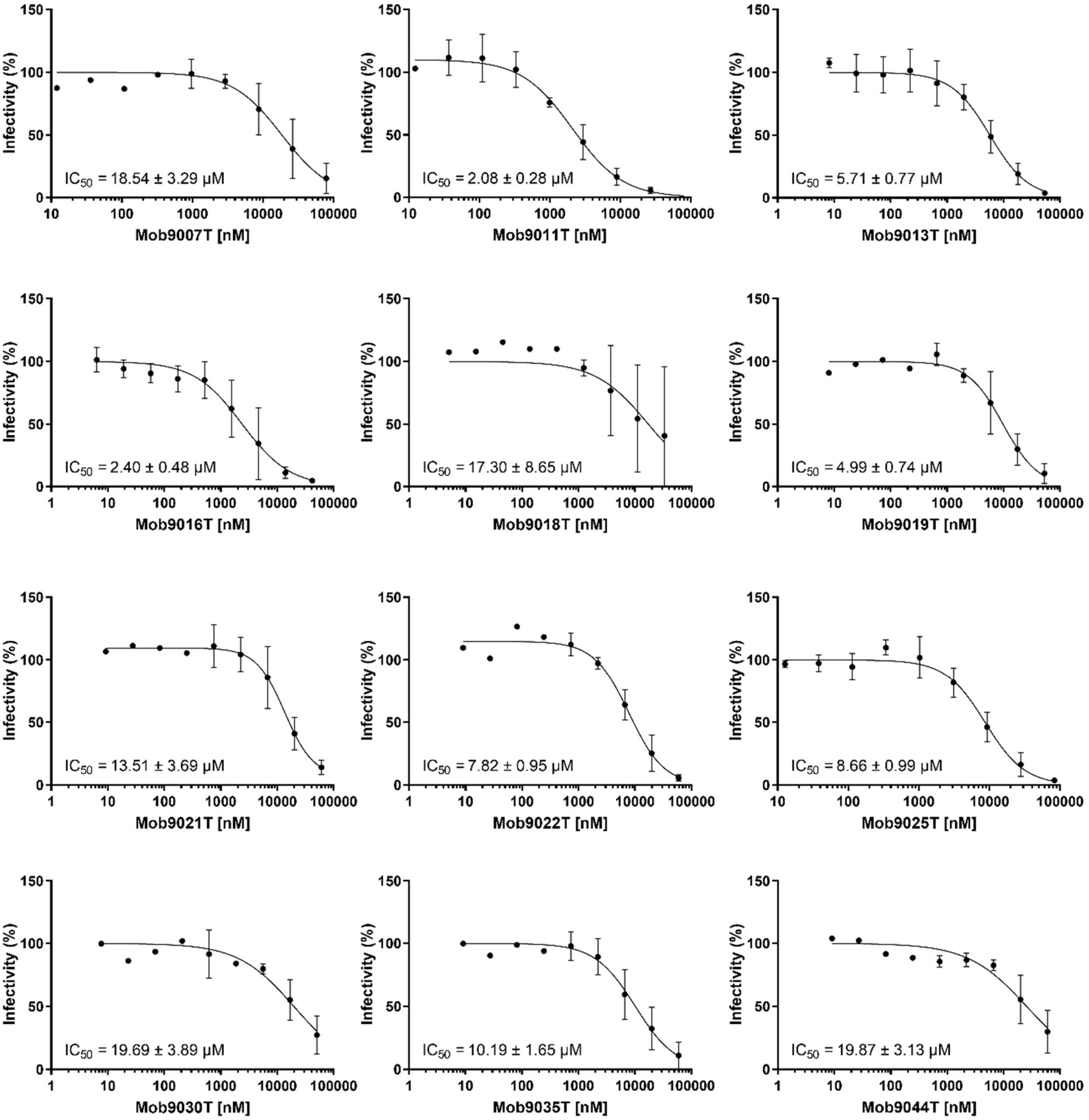


**Figure S12:** Several second-round Mobs can neutralize the SARS-CoV-2 spike pseudovirus. A dilution series of the indicated Mobs was incubated with Spike pseudotyped virus at an MOI of 0.2 for 1 hour at 37 degrees Celsius, then used to inoculate 293T-ACE2 cells in a 96-well plate. Two days later, fluorescence intensity was measured, and $IC_{50}$ values were calculated using GraphPad Prism. Data points represent the mean ± SEM of 3 independent experiments, each with 3 technical replicates.

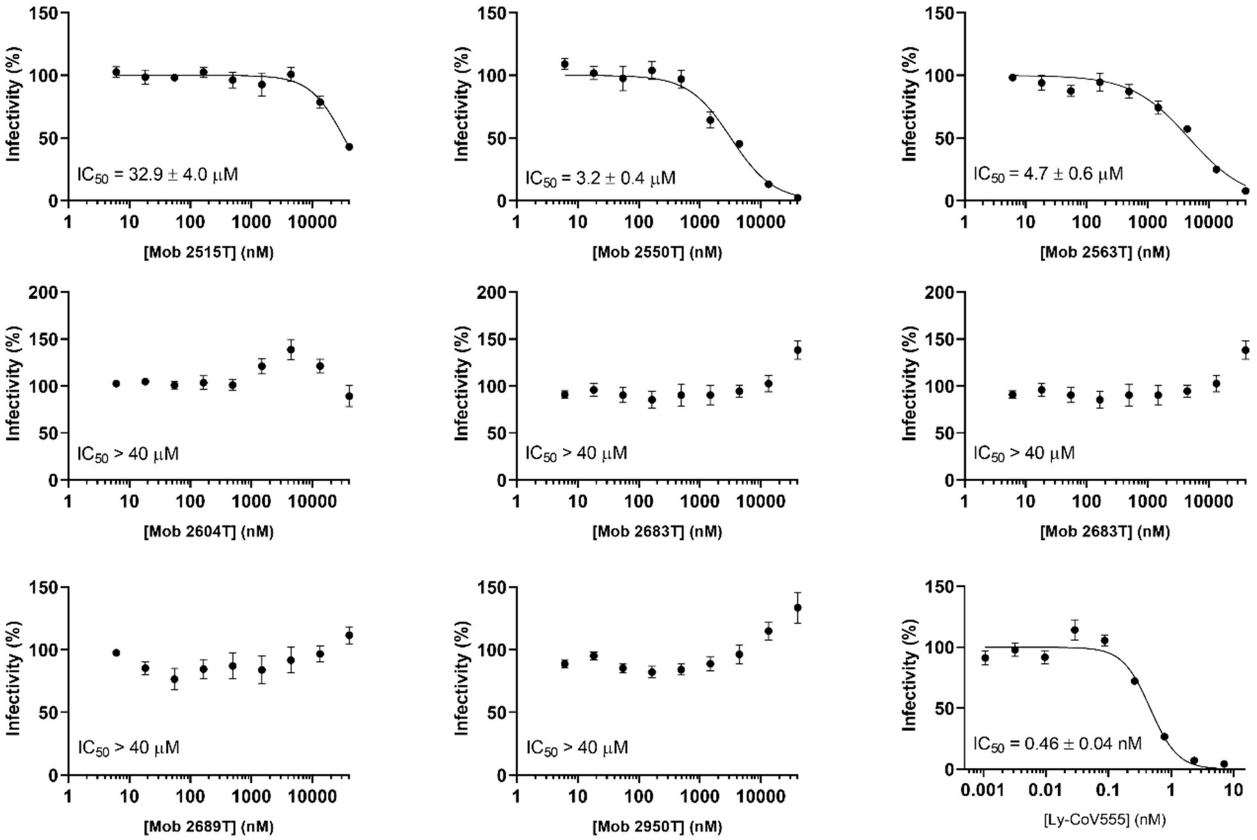
Infectivity (%)
$IC_{50}$ = 32.9 ± 4.0 μM
[Mob 2515T] (nM)
$IC_{50}$ = 3.2 ± 0.4 μM
[Mob 2550T] (nM)
$IC_{50}$ = 4.7 ± 0.6 μM
[Mob 2563T] (nM)
$IC_{50}$ > 40 μM
[Mob 2604T] (nM)
$IC_{50}$ > 40 μM
[Mob 2683T] (nM)
$IC_{50}$ > 40 μM
[Mob 2683T] (nM)
$IC_{50}$ > 40 μM
[Mob 2689T] (nM)
$IC_{50}$ > 40 μM
[Mob 2950T] (nM)
$IC_{50}$ = 0.46 ± 0.04 nM
[Ly-CoV555] (nM)

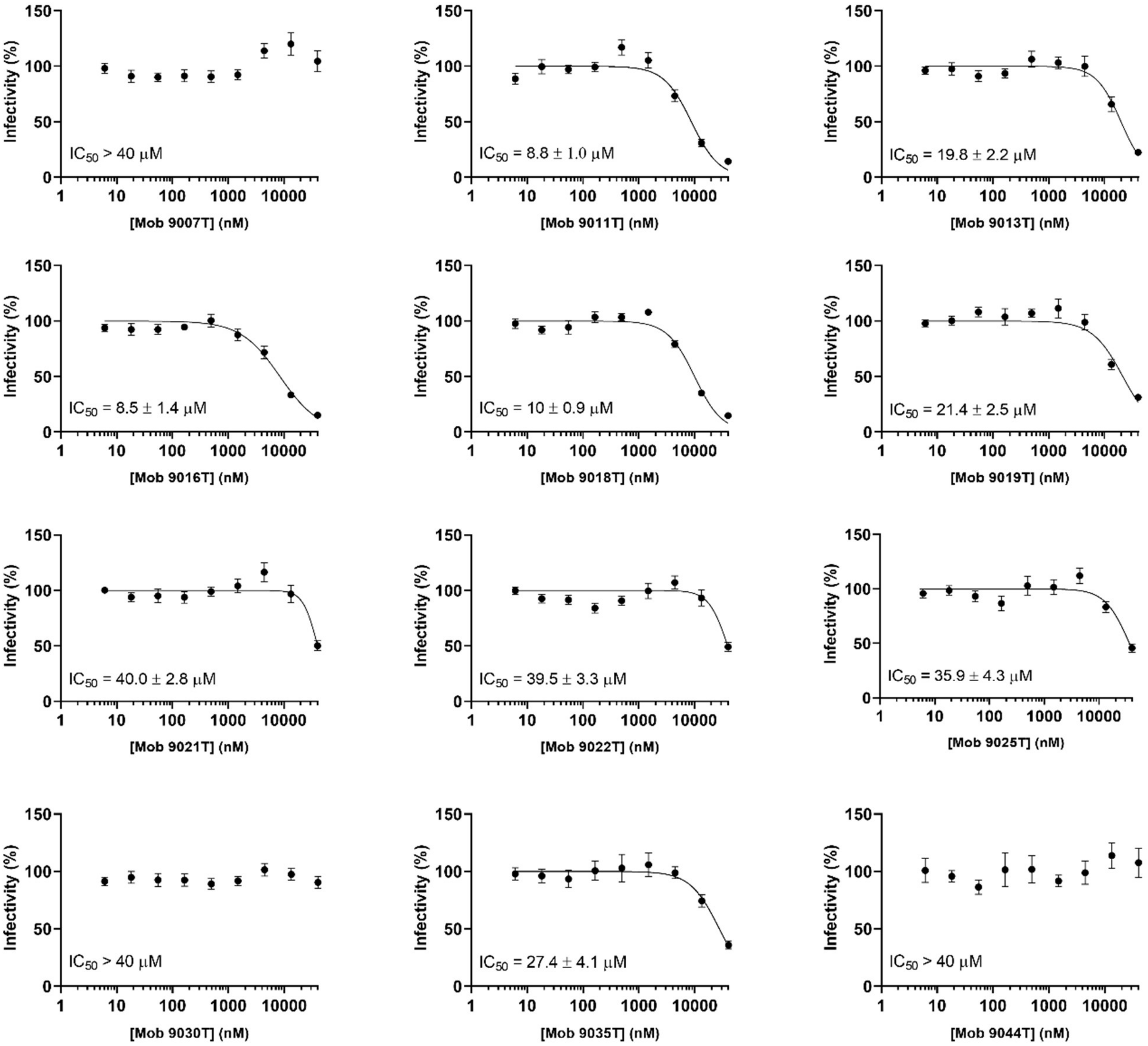


**Figure S13.** Neutralization curves and $IC_{50}$ values for Mobs against SARS-CoV-2 B.1 measured using HEK293T-ACE2-30F-PLP2 reporter cells. Threefold dilution series of each Mob were incubated with SARS-CoV-2 B.1 (MOI 0.025) before being added to cells. Firefly (FF) and Renilla (RN) activity was determined 24h after infection. FF/RN values were used to calculate the percentage of infectivity relative to maximum infectivity. Data points represent the mean ± SEM of 9 replicates pooled from three independent experiments. Human monoclonal antibody Ly-CoV555 was used as a positive control.

**Table S7:** Neutralization activity of selected Mobs against SARS-CoV-2 variants Spike-bearing pseudovirus. A dilution series of the selected Mobs was incubated with Spike pseudotyped virus at an MOI of 0.2 for 1 hour at 37 °C, then used to inoculate 293T-ACE2 cells in 96-well plates. Two days later, fluorescence intensity was measured, and $IC_{50}$ values were calculated using GraphPad Prism. Data points represent the mean ± SEM of 3 independent experiments, each with 3 technical replicates.

| Mob ID | WT (D614G) | Alpha | Delta | BA.1 | JN.1 |
|---|---|---|---|---|---|
| | $IC_{50}$ (±SEM) (μM) | $IC_{50}$ (±SEM) (μM) | $IC_{50}$ (±SEM) (μM) | $IC_{50}$ (±SEM) (μM) | $IC_{50}$ (±SEM) (μM) |
| **9011T** | **2.90** ± 0.36 | **3.51** ± 0.36 | **-** | **-** | **-** |
| **9016T** | **2.24** ± 0.28 | **4.75** ± 0.38 | **-** | **11.16** ± 1.54 | **-** |
| **2550T** | **0.79** ± 0.14 | **16.7** ± 3.0 | **0.267** ± 0.051 | **-** | **-** |
| **2563T** | **4.34** ± 0.59 | **-** | **1.221** ± 0.278 | **-** | **-** |
| Non-Mob Control **Sipavibart** | 0.420 ± 0.034 | 0.577 ± 0.044 | 0.345 ± 0.056 | 0.284 ± 0.030 | 1.96 ± 0.13 |

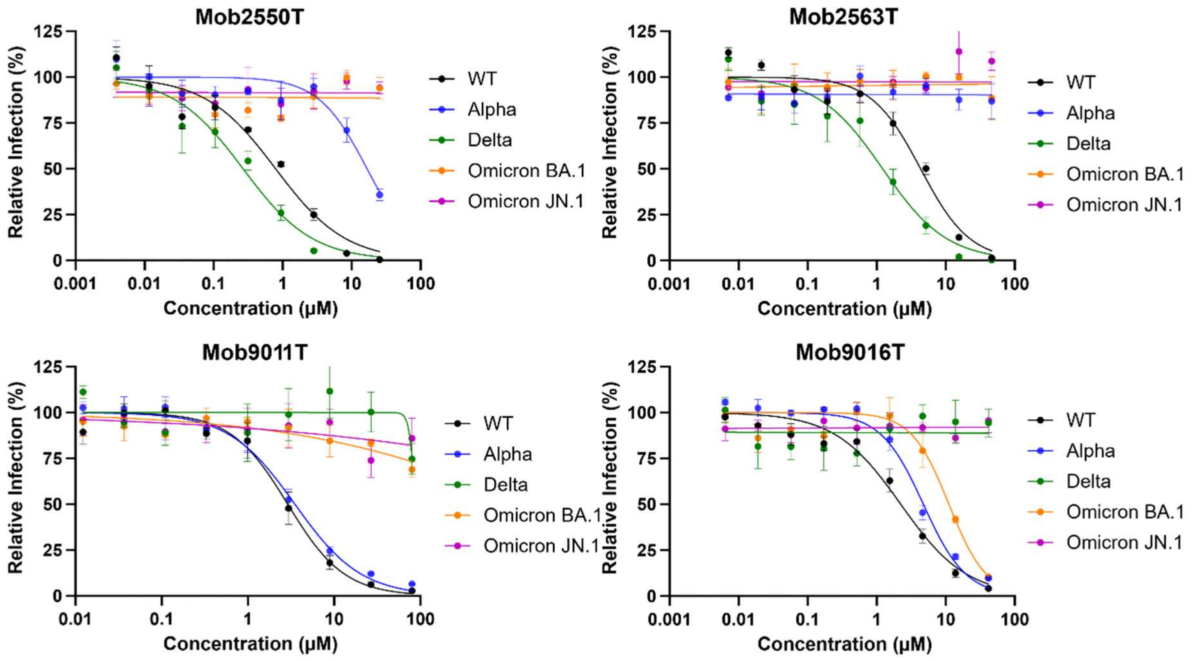


**Figure S14:** Mobs show variable neutralization of SARS-CoV-2 spike variants pseudovirus. A dilution series of the indicated Mobs was incubated with Spike pseudotyped virus at an MOI of 0.2 for 1 hour at 37 degrees Celsius, then used to inoculate 293T-ACE2 cells in a 96-well plate. Two days later, fluorescence intensity was measured, and $IC_{50}$ values were calculated using GraphPad Prism. Data points represent the mean ± SEM of 3 independent experiments, each with 3 technical replicates.

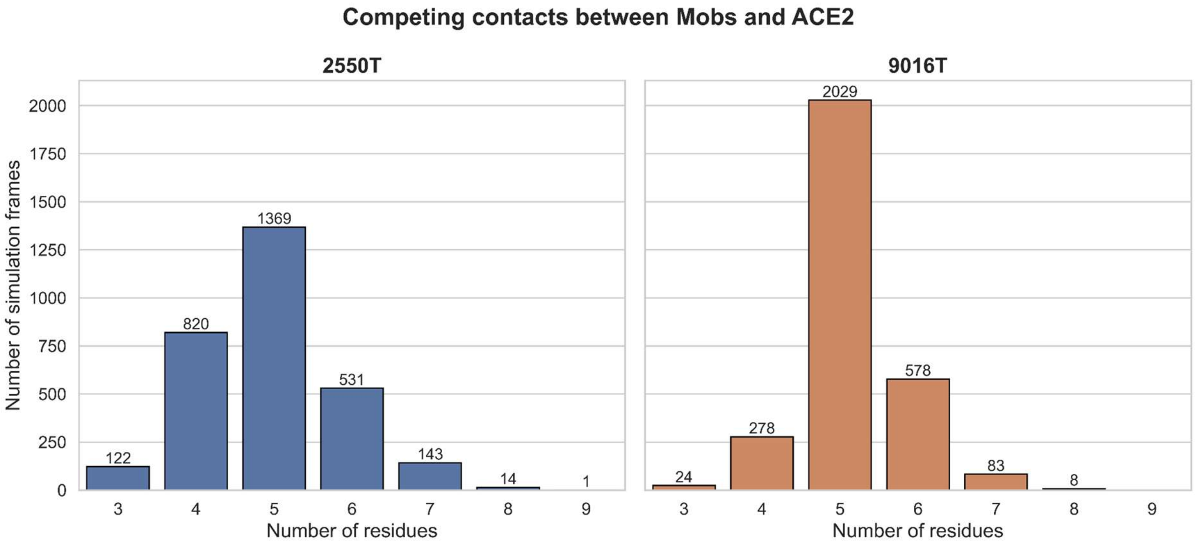


**Figure S15:** Number of residues of each simulated Mob that compete with the RBD-ACE2 interface contacts. 1000 frames were extracted from each of the 3 simulation replicates (one every 1 ns), for a total of 3000 frames per Mob. Then, through MDAnalysis (Michaud-Agrawal et al., 2011), we counted contacts in each frame. A contact was defined by a binder residue within 4 Å of the second protein. The list of RBD-ACE2 contacts was similarly compiled using PDB 6M0J.

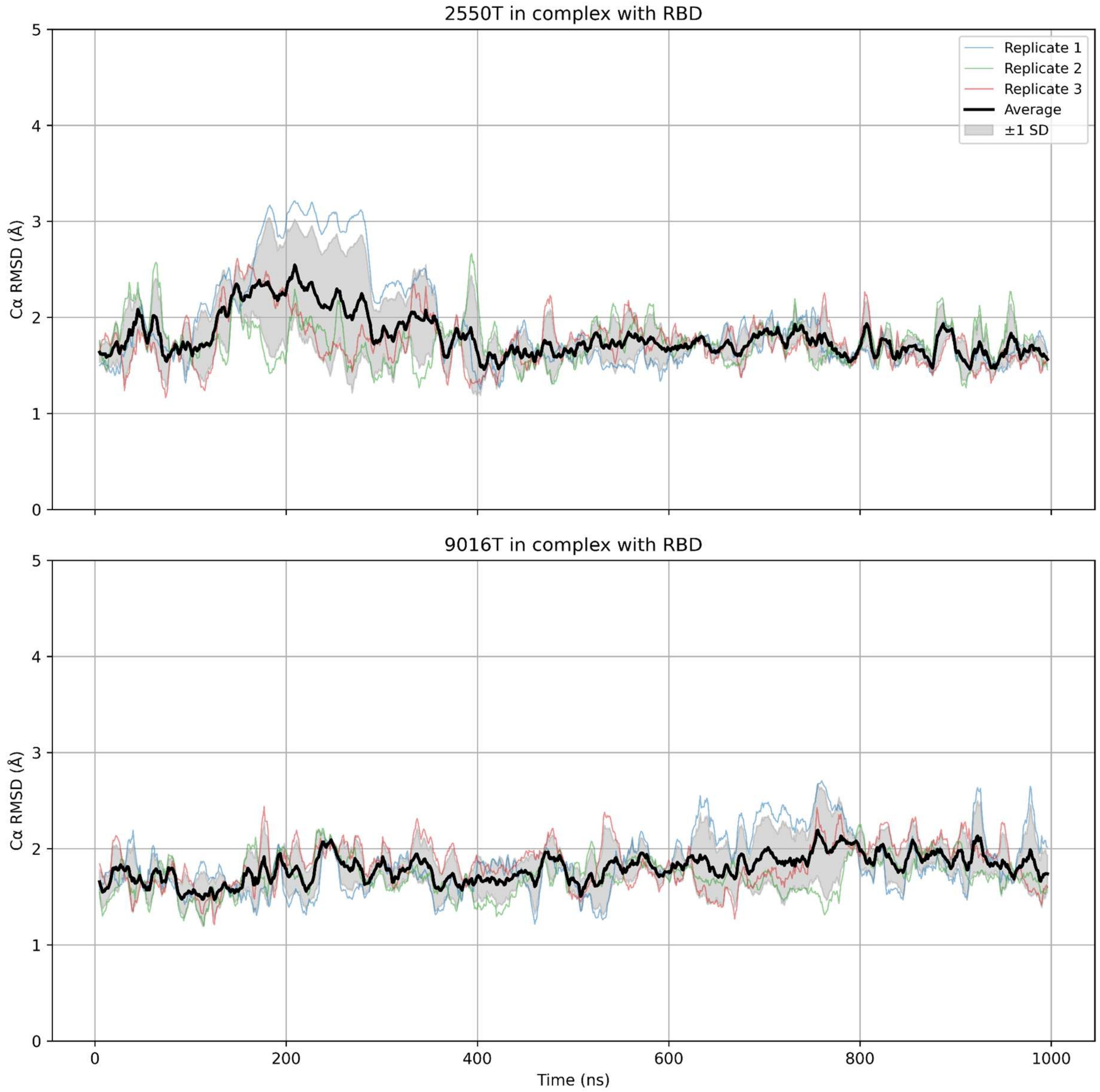


**Figure S16**: Mob RMSD of each of the two simulated neutralizing Mobs (Mob-2550T and Mob-9016T) in complex with the RBD, throughout the simulation time (1 μs), for each of the 3 replicates and the average of the replicates of the simulation. The C-terminal solubility tail is not included in the RMSD calculations.

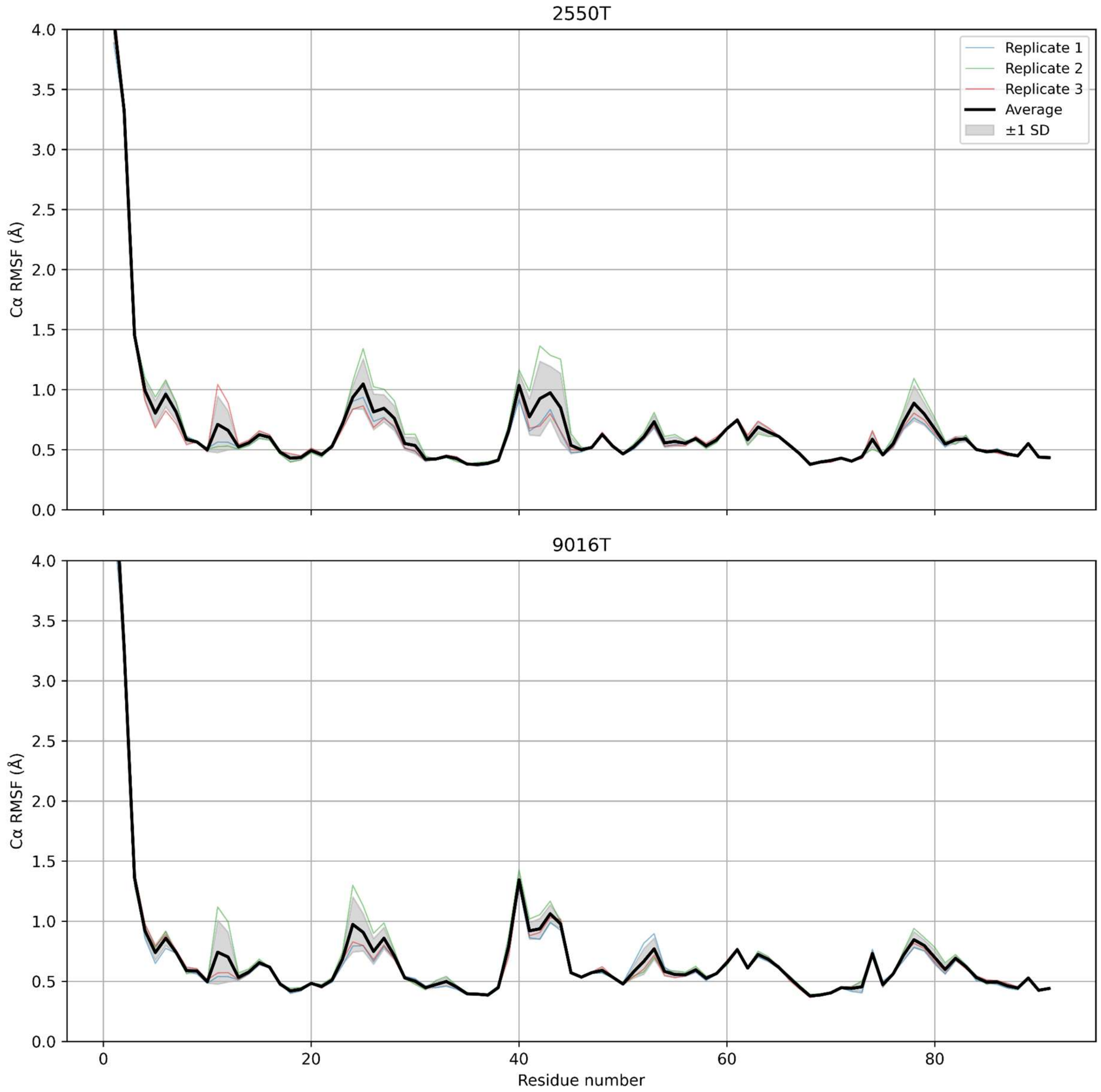


**Figure S17:** RMSF of each of the two simulated neutralizing Mobs (Mob-2550T and Mob-9016T) in complex with the RBD, throughout the simulation time, for each replicate and the average of the replicates of the simulation. The C-terminal solubility tail is not included in the RMSF calculations.

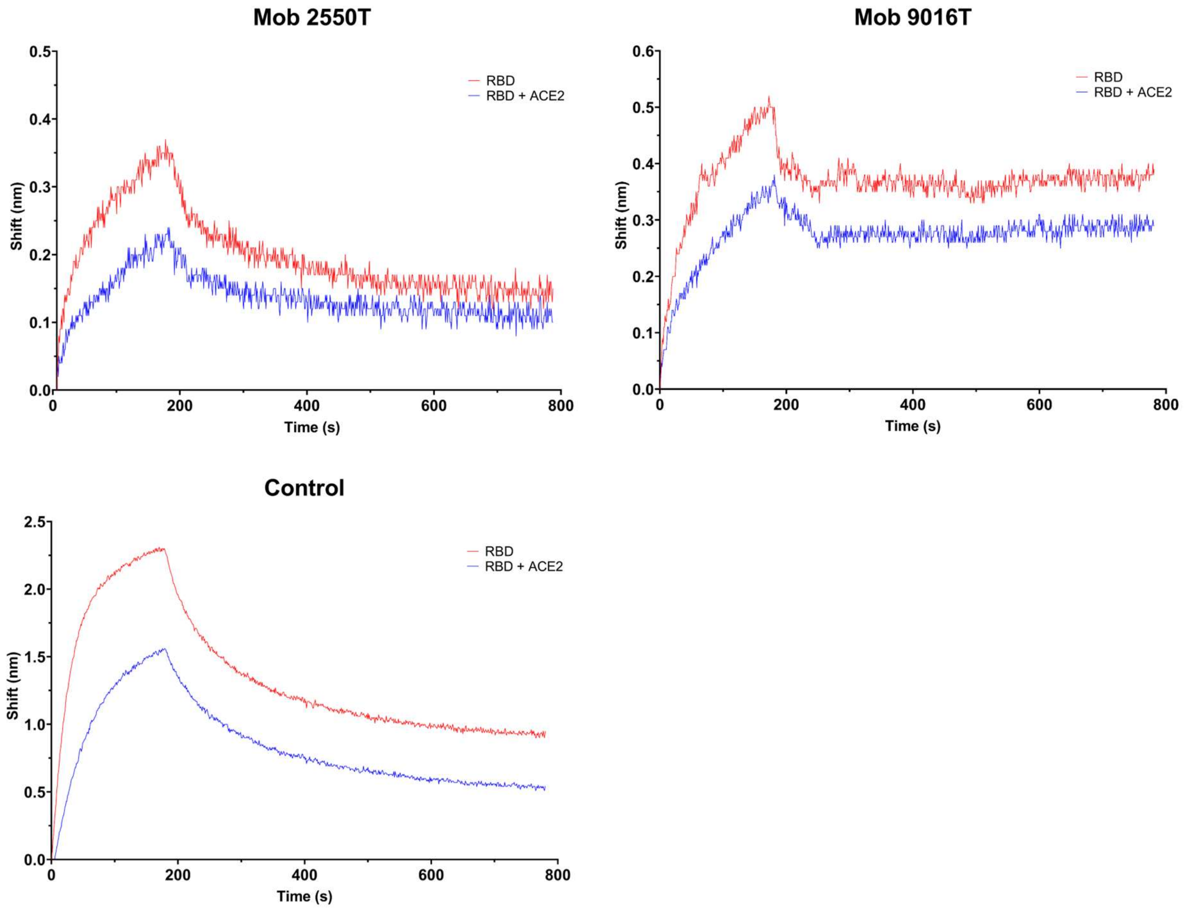


**Figure S18**: Competition assay sensorgrams of Mob 2550T and 9016T. Association and dissociation of each Mob with RBD (shown in red) and with the RBD:ACE2 complex (shown in blue). The control protein is known to bind the same epitope as ACE2.

**Table S8:** Production and characterization of spontaneous mutant Mobs arising from the YSD screening of RBD-binding Mobs, and of the same variants after addition of the C-terminal stabilizing tail (suffix "T"). Production yields were calculated as mg of pure protein (after the final size-exclusion chromatography step and the following filter sterilization) divided by the culture volume. Thermal denaturation and aggregation were analyzed with a Nanotemper Prometheus Panta, thereby estimating the melting temperature ($T_m$) by DSF, and the onset of aggregation temperature for large insoluble aggregates ($T_{on\_turbidity}$, obtained from light backscattering turbidometry) and/or soluble aggregates ($T_{on\_scattering}$ , obtained by DLS).

| Mob ID | Yield (mg/L culture) | $T_m$ (°C) | $T_{on\_turbidity}$ (°C) | $T_{on\_scattering}$ (°C) |
|---|---|---|---|---|
| **3560** | 13 | 47.3 ± 0.1 | no | n.d. |
| **3561** | 10 | 47.7 ± 0.4 | no | 45.7 ± 2.0 |
| **3562** | 10 | 48.1 ± 0.5 | no | n.d. |
| **3563** | 25 | 46.1 ± 0.2 | no | n.d. |
| **3564** | 33 | 48.2 ± 0.9 | no | n.d. |
| **3565** | 7 | 43.3 ± 0.1 | no | 41.3 ± 2.1 |
| **3566** | 7 | 43.3 ± 0.3 | no | 35.2 ± 0.6 |
| **3567** | 15 | 45.7 ± 0.1 | no | 54.9 ± 0.6 |
| **3569** | 23 | 47.7 ± 0.1 | no | n.d. |
| **3560T** | 94 | 60.6 ± 0.1 | no | 54.2 ± 0.2 |
| **3561T** | 72 | 57.5 ± 0.1 | no | 51.6 ± 1.2 |
| **3563T** | 176 | 57.0 ± 0.1 | no | 50.7 ± 0.3 |

**Table S9:** Affinity for RBD (**Figure S19**) and pseudovirus & virus neutralization (**Figures S20** and **S21**, respectively) capacity of YSD-discovered mutant Mobs after addition of stabilizing C-terminal tail.

| **Mob ID** | **$k_{on}$ (± SE) ($M^{-1}$·$s^{-1}$)** | **$k_{off}$ (± SE) ($s^{-1}$)** | **$K_D$ (± SE) (nM)** | **$IC_{50}$ (± SEM) (µM) (pseudovirus)** | **$IC_{50}$ (± SEM) (µM) (virus)** |
|---|---|---|---|---|---|
| **3560T** | 4.88 ± 3.53 E05 | 1.91 ± 0.64 E-02 | 39.1 ± 31.2 | 9.54 ± 2.17 | 33.8 ± 4.8 |
| **3561T** | 1.15 ± 0.24 E05 | 4.52 ± 1.20 E-03 | 39.4 ± 13.3 | >25.75 | 25.8 ± 3.1 |
| **3563T** | 1.92 ± 0.22 E05 | 8.10 + 1.17 E-03 | 42.3 ± 7.9 | 6.19 ± 0.80 | 26.7 ± 3.2 |

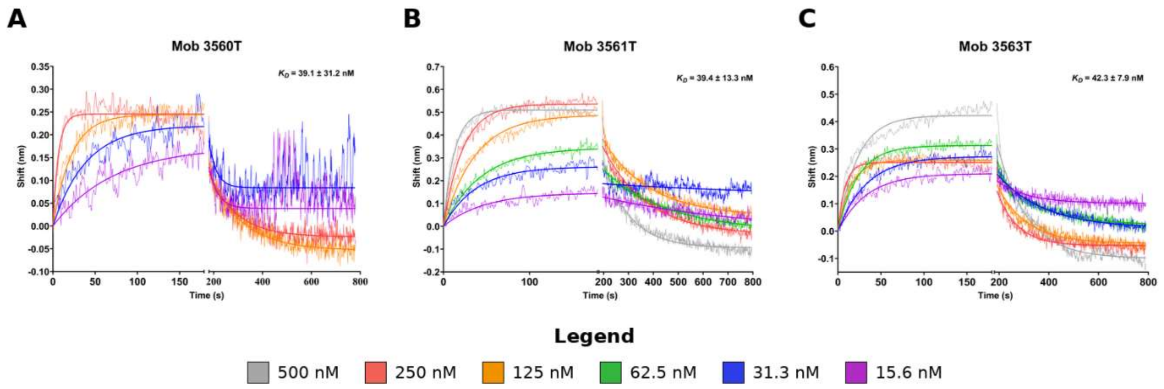


**Figure S19:** SPR sensorgrams for each of YSD-discovered mutant Mobs after addition of stabilizing C-terminal tail with measurable $K_D$ values. Association and dissociation curves were measured at different RBD concentrations for each tested Mob.

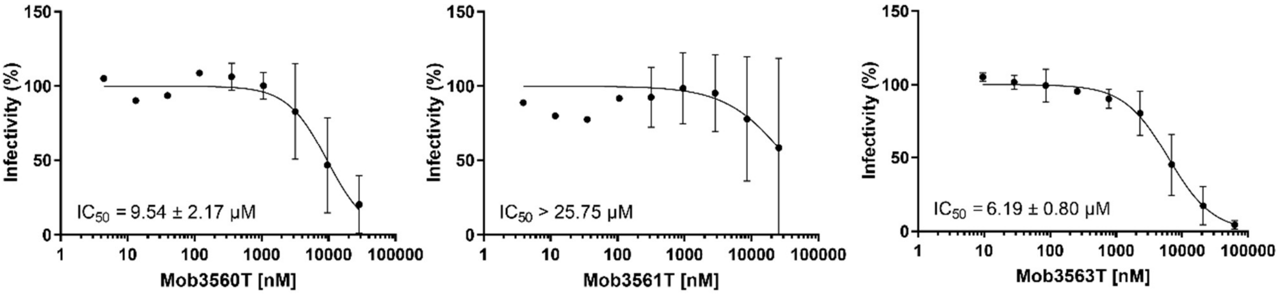


**Figure S20:** Some of the YSD-discovered mutant Mobs show variable neutralization of SARS-CoV-2 spike variants pseudovirus. A dilution series of the indicated Mobs was incubated with Spike pseudotyped virus at an MOI of 0.2 for 1 hour at 37 degrees Celsius, then used to inoculate 293T-ACE2 cells in a 96-well plate. Two days later, fluorescence intensity was measured, and $IC_{50}$ values were calculated using GraphPad Prism. Data points represent the mean ± SEM of 3 independent experiments, each with 3 technical replicates.

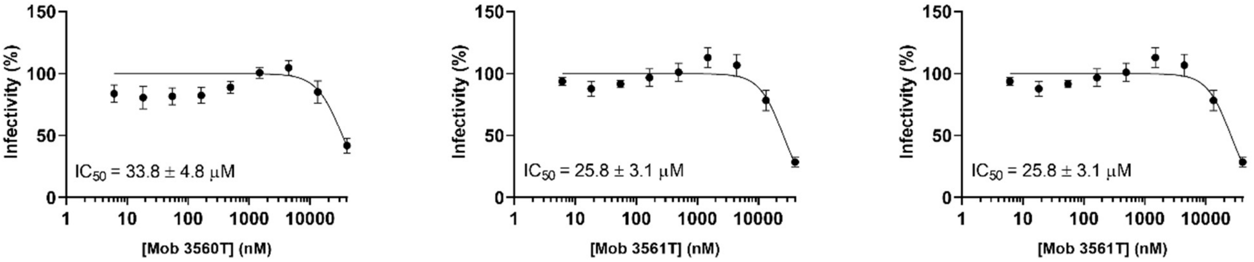


**Figure S21.** Neutralization curves and $IC_{50}$ values for YSD-discovered mutant Mobs against SARS-CoV-2 B.1 measured using HEK293T-ACE2-30F-PLP2 reporter cells. Threefold dilution series of each Mob were incubated with SARS-CoV-2 B.1 (MOI 0.025) before being added to cells. Firefly (FF) and Renilla (RN) activity was determined 24h after infection. FF/RN values were used to calculate the percentage of infectivity relative to maximum infectivity. Data points represent the mean ± SEM of 3 replicates pooled from three independent experiments. Human monoclonal antibody Ly-CoV555 was used as a positive control.

**Table S10.** Sequences of the expressed Mobs. Bold amino acids correspond to the designed Mob sequences. The suffix "(T)" in the Mob ID indicates that the Mob was expressed both with and without the C-terminus solubility tail; "T" indicates it was expressed with the tail only.

| Mob | Sequence |
|---|---|
| 2515(T) | MHHHHHHSGTSG**VSDVPRDLEVVAATPTSLLISWDASTPDVAVYMIVYHETKDPNNAMQFTVPHNISTATILGLQPGVEYTIYVLAIMADGTVPDAIEINHRT**(EIDKPSQ) |
| 2550(T) | MHHHHHHSGTSG**VSDVPRDLEVVAATPTSLLISWDASIPNVVRYLIVYHPTGYYSEWQQFAVPGTAAYARIQGLQPGVEYTIYVLGILADGTVLDQHEINGKT**(EIDKPSQ) |
| 2563(T) | MHHHHHHSGTSG**VSDVPRDLEVVAATPTSLLISWDASIPNVVRYLIVYHPTGYDSEWQQKAVPGTAAYARIQGLQPGVEYTIKVLGILADGTVLDEHEINGKT**(EIDKPSQ) |
| 2604(T) | MHHHHHHSGTSG**VSDVPRDLEVVAATPTSLLISWDASIPNVVRYLIIYHPTGYYSEYQQKAVPGTAAYARIQGLQPGVEYTIFVLGILADGTVLEQHEINGKT**(EIDKPSQ) |
| 2683(T) | MHHHHHHSGTSG**VSDVPRDLEVVAATPTSLLISWDASIPNVVKYLIVYHPTGYYSEWQQFAVDGTAAYARIQGLQPGVEYTIKVLGILADGTVLDEHEINGKT**(EIDKPSQ) |
| 2689(T) | MHHHHHHSGTSG**VSDVPRDLEVVAATPTSLLISWDATPPASERVLFILIIYGEAGGNTPFQSFTVPPSLRNATISGLKAGVEYTIHVLAATVTSNAPDAPVDVNQISINATT**(EIDKPSQ) |
| 2950(T) | MHHHHHHSGTSG**VSDVPRDLEVVAATPTSLLISWDASIPNVVRYLIIYHPTGYYSEFQQFAVPGTAAYARIQGLQPGVEYTIFVLGILADGTVLDEHEINGKT**(EIDKPSQ) |
| 3560(T) | MHHHHHHSGTSG**VSDVPRDLEVVAATPTSLLISWDASIPNVVKYLIVYHPTGYYSEWQQFAVPGTAAYARIQGLQPGVEYTIYVLGILADGTVLDQHEINGKT**(EIDKPSQ) |
| 3561(T) | MHHHHHHSGTSG**VSDVPRDLEVVAATPTSLLISWDASIPNVVRYLIIYHPTGYYSEWQQFAVPGTAAYARIQGLQPGVEYTIYVLGILADGTVLDQHEINGKT**(EIDKPSQ) |
| 3562 | MHHHHHHSGTSG**VSDVPRDLEVVAATPTSLLISWDASIPNVVRYLIVYHPTGYYSEWQQFAVPGTAAYARIQGLQPGVEYTIYVLGILADGTVLDQHEINDKT** |
| 3563(T) | MHHHHHHSGTSG**VSDVPRDLEVVAATPTSLLISWDASIPNVVKYLIVYHPTGYYSEWQQFAVDGTAAYARIQGLQPGVEYTIYVLGILADGTVLDQHEINGKT**(EIDKPSQ) |
| 3564 | MHHHHHHSGTSG**VSDVPRDLEVVAATPTSLLISWDASIPNVVKYLIVYHPTGYYSEWQQFAVDGTAAYARIQGLQPGVEYTIYVLGILADGTVLDQHEINDKT** |
| 3565 | MHHHHHHSGTSG**VSDVPRDLEVVAATPTSLLISWDASIPNVVRYLIIYHPTGYYSEFQQFAVPGTAAYARIQGLQPGVEYTIFVLGILADGTVLDQHEINGKT** |
| 3566 | MHHHHHHSGTSG**VSDVPRDLEVVAATPTSLLISWDASIPNVVRYLIIYHPTGYYSEFQQFAVPGTAAYARIQGLQPGVEYTIFVLGILADGTVLEQHEINGKT** |
| 3567 | MHHHHHHSGTSG**VSDVPRDLEVVAATPTSLLISWDASIPNVVRYLIIYHPTGYYSEFQQFAVPGTAAYARIQGLQPGVEYTIFVLGILADGTVLEQHEINDKT** |
| 3569 | MHHHHHHSGTSG**VSDVPRDLEVVAATPTSLLISWDATPPASERVLFILIIYGEAGGNTPFQSFTVPPSLRNATISGLKAGVEYTIHVLAATVTSNAPDAPVDVNEISINATT** |
| 9007T | MHHHHHHSGTSG**VSDVPRDLEVVAATPTSLLISWDASIPNVVTYLIVYHPTGYDSEWQQMAVPGTAAYARIQGLQPGVEYTIKVLGILADGTVLDEHTINGKT**EIDKPSQ |
| 9011T | MHHHHHHSGTSG**VSDVPRDLEVVAATPTSLLISWDASIPNVVTFLIIYHPTGRTSEWQQFAVPGTAAYARIQGLQPGVEYTIYVLGILADGEVLDQHEINGKT**EIDKPSQ |
| 9013T | MHHHHHHSGTSG**VSDVPRDLEVVAATPTSLLISWDASIPNVVTYLIVYHPTGRYSEWQQFAVPGTAAYARIQGLQPGVEYTIYVLGILEDGTVMDQHEINGKT**EIDKPSQ |
| 9016T | MHHHHHHSGTSG**VSDVPRDLEVVAATPTSLLISWDASIPNVVAYLIVYHPTGYDSEWQQFAVPGTAAYARIQGLQPGVEYTIYVLGIQADGTVLDQHEINGKT**EIDKPSQ |
| 9018T | MHHHHHHSGTSG**VSDVPRDLEVVAATPTSLLISWDASIPNVVSYLIVYHPTGYYSEWQQFAVPGTAAYARIQGLQPGVEYQIYVLGILADGTVMDQHEINGKT**EIDKPSQ |
| 9019T | MHHHHHHSGTSG**VSDVPRDLEVVAATPTSLLISWDASIPNVVTYLIVYHPTGKESEWQQFAVPGTAAYARIQGLQPGVEYTIYVLGILADGTVLDQHEINGKT**EIDKPSQ |
| 9021T | MHHHHHHSGTSG**VSDVPRDLEVVAATPTSLLISWDASIPNVVRYLIVYYPTGYYDEWQQFALPGTAAYARIQGLQPGVEYTIYVLGILADGTVLDQHEINGKT**EIDKPSQ |
| 9022T | MHHHHHHSGTSG**VSDVPRDLEVVAATPTSLLISWDASIPNVVTYLIVYHPTGYESEWQQFAVPGTAAYARIQGLQPGVEYTIYVLGILADGAVLDQHEINGKT**EIDKPSQ |
| 9025T | MHHHHHHSGTSG**VSDVPRDLEVVAATPTSLLISWDASIPNVVTYLIVYHPTGYESEWQQFAVPGTAAYARIQGLQPGVEYTIYVLGILADGTVLEQHEINGKT**EIDKPSQ |
| 9030T | MHHHHHHSGTSG**VSDVPRDLEVVAATPTSLLISWDASVPNVAVYMVVYHATDDPNNAKQFTVPPTQSSALITGLEAGKTYRIYVLAIMADGSVPDAAEILATT**EIDKPSQ |
| 9035T | MHHHHHHSGTSG**VSDVPRDLEVVAATPTSLLISWDASVPDVAVYMIVYHETTDPNDAKQFTVPATKSTAEIQGLKSGVTYTVYVLAIMADGTVPDAAELQATT**EIDKPSQ |
| 9044T | MHHHHHHSGTSG**VSDVPRDLEVVAATPTSLLISWDASVPNVARFMIVYHEESDPNSAQQFTVPASESTALISGLTPGKKYTIYVLAILADGTVPDAMEITATT**EIDKPSQ |
